# The quorum-sensing systems of *Vibrio campbellii* DS40M4 and BB120 are genetically and functionally distinct

**DOI:** 10.1101/2020.03.31.019307

**Authors:** Chelsea A. Simpson, Blake D. Petersen, Nicholas W. Haas, Logan J. Geyman, Aimee H. Lee, Ram Podicheti, Robert Pepin, Laura C. Brown, Douglas B. Rusch, Michael P. Manzella, Kai Papenfort, Julia C. van Kessel

**Affiliations:** Department of Biology, Indiana University, Bloomington, IN 47405; Center for Genomics and Bioinformatics, Indiana University, Bloomington, IN 47405; Mass Spectrometry Facility, Indiana University, Bloomington, IN 47405; Department of Chemistry, Indiana University, Bloomington, IN 47405; Friedrich Schiller University, Institute of Microbiology, 07745 Jena, Germany; Microverse Cluster, Friedrich Schiller University Jena, 07743 Jena, Germany

**Keywords:** quorum sensing, *Vibrio*, *Vibrio campbellii*, natural transformation, biofilm, protease

## Abstract

*Vibrio campbellii* BB120 (previously classified as *Vibrio harveyi*) is a fundamental model strain for studying quorum sensing in vibrios. A phylogenetic evaluation of sequenced *Vibrio* strains in Genbank revealed that BB120 is closely related to the environmental isolate *V. campbellii* DS40M4. We exploited DS40M4’s competence for exogenous DNA uptake to rapidly generate >30 isogenic strains with deletions of genes encoding BB120 quorum-sensing system homologs. Our results show that the quorum-sensing circuit of DS40M4 is distinct from BB120 in three ways: 1) DS40M4 does not produce an acyl homoserine lactone autoinducer but encodes an active orphan LuxN receptor, 2) the quorum regulatory small RNAs (Qrrs) are not solely regulated by autoinducer signaling through the response regulator LuxO, and 3) the DS40M4 quorum-sensing regulon is much smaller than BB120 (~100 genes vs ~400 genes, respectively). Using comparative genomics to expand our understanding of quorum-sensing circuit diversity, we observe that conservation of LuxM/LuxN proteins differs widely both between and within *Vibrio* species. These strains are also phenotypically distinct: DS40M4 exhibits stronger interbacterial cell killing, whereas BB120 forms more robust biofilms and is bioluminescent. These results underscore the need to examine wild isolates for a broader view of bacterial diversity in the marine ecosystem.

**Originality-Significance Statement:** Wild bacterial isolates yield important information about traits that vary within species. Here, we compare environmental isolate *Vibrio campbellii* DS40M4 to its close relative, the model strain BB120 that has been a fundamental strain for studying quorum sensing for >30 years. We examine several phenotypes that define this species, including quorum sensing, bioluminescence, and biofilm formation. Importantly, DS40M4 is naturally transformable with exogenous DNA, which allows for the rapid generation of mutants in a laboratory setting. By exploiting natural transformation, we genetically dissected the functions of BB120 quorum-sensing system homologs in the DS40M4 strain, including two-component signaling systems, transcriptional regulators, and small RNAs.

## Introduction

Quorum sensing is a form of cell-cell communication between bacteria in which individual cells synthesize and respond to signaling molecules called autoinducers. Increasing concentrations of autoinducers trigger changes in gene expression among bacterial populations, resulting in group behaviors. Through this method of cell-cell signaling, bacteria control expression of genes involved in motility, toxin secretion, metabolism, and more (Rutherford and Bassler, 2012). Quorum sensing has been heavily studied in marine *Vibrio* species, owing to easily observable behaviors such as bioluminescence and biofilm formation (Bassler *et al*., 1997). *Vibrio campbellii* BB120 was one of the first *Vibrio* species to have its quorum-sensing circuit discovered (Bassler *et al*., 1993, 1994; Freeman *et al*., 2000). *V. campbellii* BB120 was historically called *Vibrio harveyi* BB120 (also known as strain ATCC BAA-1116) until it was recently reclassified (Lin *et al*., 2010). Although similar quorum-sensing system architectures have been established in other vibrios, BB120 has remained a model organism for studying quorum sensing and signal transduction among the *Vibrionaceae*.

In the *V. campbellii* BB120 quorum-sensing system, previous research has identified three membrane-bound histidine kinase receptors, CqsS, LuxPQ, and LuxN (Fig. 1A) (J. M. Henke and Bassler, 2004; Rutherford and Bassler, 2012; Jung *et al*., 2016). These receptors bind to autoinducers CAI-1 ((*S*)-3-hydroxytridecan-4-one), AI-2 ((2*S*, 4*S*)-2-methyl-2,3,4-tetrahydroxytetrahydrofuran-borate), and HAI-1 (N-(3-hydroxybutyryl)-homoserine lactone), respectively. These autoinducers are constitutively produced by the autoinducer synthases, CqsA, LuxS, and LuxM, respectively. A cytosolic hybrid histidine kinase, termed HqsK (also called VpsS in *Vibrio cholerae*) that interacts with a nitric oxide (NO) sensing protein (H-NOX) has also been identified as part of this pathway (Henares *et al*., 2012; Jung *et al*., 2015). When cells are at low cell density (LCD), the local concentration of autoinducers produced by *V. campbellii* is insufficient to bind to the receptors, which results in the receptors functioning as kinases. In the absence of NO, HqsK also acts as a kinase. In this LCD state, all four receptors phosphorylate LuxU, a phosphotransfer protein that transfers phosphate to the response regulator LuxO (Fig. 1). Phosphorylated LuxO activates transcription of five small RNAs (sRNAs) termed quorum regulatory RNAs (Qrrs). The Qrrs post-transcriptionally repress production of the transcriptional regulator LuxR while activating expression of the transcriptional regulator AphA. The activation of AphA and repression of LuxR combinatorially regulates ~170 genes and produces individual behaviors in *V. campbellii*, including biofilm formation (Fig. 1) (van Kessel *et al*., 2013). As *V. campbellii* cells continue to grow and the local concentration of autoinducers produced by cells increases, the autoinducers bind to their cognate receptors and change their kinase activity to act as phosphatases. Likewise, in the presence of NO, the H-NOX protein binds NO, inhibiting the kinase activity of HqsK. In this high cell density (HCD) state, LuxO is not phosphorylated, and the Qrrs are no longer expressed. The result is high expression of LuxR and no production of AphA. Based on transcriptomic data, LuxR regulates >400 genes to produce group behaviors in *V. campbellii*, including activation of bioluminescence, type VI secretion (T6SS), and protease production, and repression of biofilm production and type III secretion (T3SS, Fig. 1) (van Kessel *et al*., 2013; Bagert *et al*., 2016).

**Figure 1:**
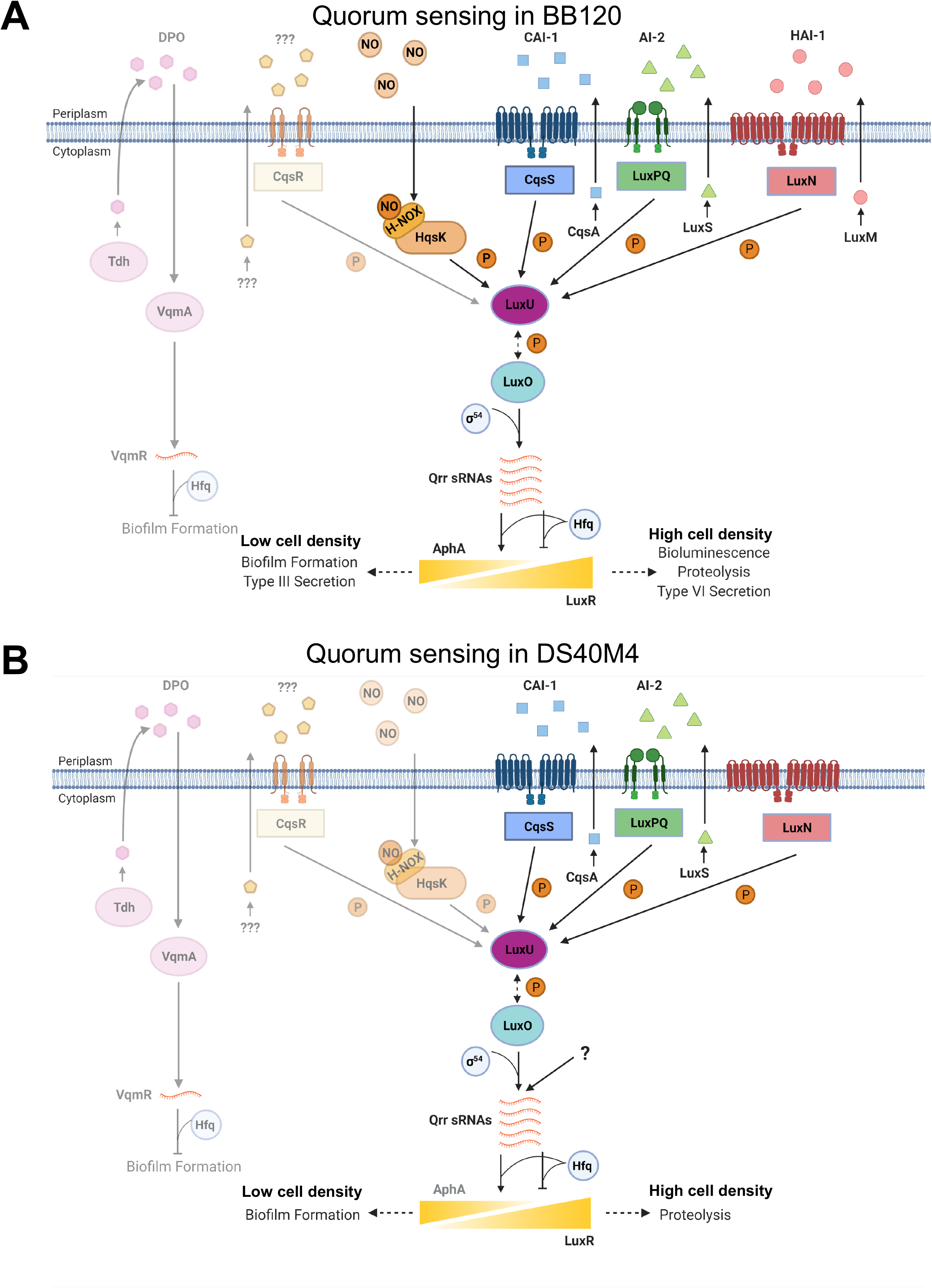
Model for quorum-sensing regulation in *V. campbellii* strains BB120 and DS40M4. The quorum-sensing models in *V. campbellii* BB120 (A) and DS40M4 (B). Darker colored circuits represent pathways that have been tested in each strain. Lighter colored circuits include putative homologs to other *Vibrio* systems that have not yet been tested in these strains. (A) In the established BB120 pathway, the autoinducers CAI-1, HAI-1, AI-2, are produced by autoinducer synthases CqsA, LuxM, and LuxS and sensed by receptors CqsS, LuxN, and LuxPQ, respectively. At LCD, autoinducers are at low concentrations, resulting in the receptors acting as kinases. The receptors phosphorylate LuxU (phosphorelay protein), which transfers the phosphate to LuxO. Phosphorylated LuxO activates *qrr* expression through Sigma-54. The Qrrs together with Hfq bind to the *aphA* and *luxR* mRNAs, and AphA is expressed and LuxR production is minimal. The combination of AphA and LuxR protein levels leads to LCD behaviors, such as T3SS and biofilm formation. As autoinducers accumulate at HCD, the receptors bind autoinducers and in this state act as phosphatases. De-phosphorylated LuxO does not activate the *qrr* genes, thus leading to maximal LuxR production and absence of AphA. This ultimately leads to HCD behaviors such as bioluminescence, proteolysis, and type VI secretion (T6SS). In the absence of nitric oxide (NO), HqsK acts as a kinase. In the presence of NO, NO-bound H-NOX inhibits the kinase activity of HqsK contributing to the de-phosphorylation of LuxO. In *V. cholerae*, the CqsR receptor senses an unknown autoinducer from an unidentified synthase. In the *V. cholerae* DPO-dependent QS pathway, the autoinducer DPO is produced from threonine catabolism which is dependent on the Tdh enzyme. Extracellular DPO binds to the receptor VqmA. The VqmA-DPO complex activates transcription of the VqmR sRNA, which represses genes required for biofilm formation and toxin production. (B) Our proposed model for quorum-sensing regulation in *V. campbellii* DS40M4. The CqsA/CqsS and LuxS/LuxPQ receptor pairs are conserved and act as described above. The LuxN receptor is active either in the absence of a ligand or in the presence of an unidentified ligand other than an AHL, but the mechanism by which it functions is undetermined. We propose that there is an additional regulator outside of the LuxO pathway that also converges on the Qrrs. At HCD, LuxR controls fewer behaviors than in BB120. Image created with BioRender.com.

Recently, we showed that natural transformation is achievable in two wild isolates of *V. campbellii*, DS40M4 and NBRC 15631, by ectopically inducing expression of the *V. cholerae* competence regulator TfoX (Simpson *et al*., 2019). By exploiting this process, researchers are able to make marked and unmarked gene deletions, point mutations, and insertions using transformation (Meibom *et al*., 2005; Yamamoto *et al*., 2011; Sun *et al*., 2013). Importantly, multiple mutations can be generated simultaneously in *V. cholerae* through a technique termed MuGENT: multiplex genome editing by natural transformation (Dalia *et al*., 2017; Dalia, 2018). While *V. campbellii* strains all contain homologs of the known competence genes, TfoX overexpression does not stimulate expression of these genes in BB120, unlike in the wild isolates DS40M4 and NBRC 15631 (Simpson *et al*., 2019). This finding suggested that BB120 may have lost regulation by TfoX due to domestication in laboratory conditions. We hypothesized that environmental *V. campbellii* strains may exhibit differences in other behaviors such as growth and quorum sensing-controlled phenotypes. Using natural transformation, we systematically constructed isogenic mutant strains of DS40M4 with deletions in predicted quorum-sensing components. With these mutants, we assessed the regulatory circuit of DS40M4 compared to BB120. In addition, we tested differences in quorum sensing-controlled phenotypes between DS40M4 and BB120, including biofilm formation, interbacterial cell killing, protease activity, and bioluminescence. Altogether, our data show that while DS40M4 is similar to BB120 in the presence of specific genes, it has a distinct quorum-sensing circuit that produces different downstream behaviors.

## Results

### V. campbellii DS40M4 has an active quorum-sensing system

We had previously shown that DS40M4 encodes a LuxR and LuxO homolog that function to regulate bioluminescence similarly to the epistasis observed in BB120 (Simpson *et al*., 2019). We have also shown that DS40M4 responds to autoinducers AI-2 and CAI-1 produced by BB120. However, we had not examined the presence and function of other core quorum-sensing components. DS40M4 is closely related to BB120; for example, chromosome I of both organisms share 97.45% nucleotide identity. Thus, we predicted that much of the quorum sensing circuit would be conserved in DS40M4. As a comparison, most of the BB120 quorum sensing circuit architecture is conserved in *V. parahaemolyticus*, which is more distantly related to BB120 (chromosome I of both organisms share 83.58% nucleotide identity) (Bassler *et al*., 1997; J. Henke and Bassler, 2004; Gode-Potratz and McCarter, 2011). Thus, we searched the DS40M4 genome for homologs of the proteins and sRNAs previously identified in the *V. campbellii* BB120 quorum-sensing system. DS40M4 encodes genes with high amino acid or nucleotide identity to all the components except for LuxM, for which there was no homolog in DS40M4 (Table 1). We also identified homologs of the HqsK/VpsS and H-NOX system from BB120 (Fig. 1, Table 1). There have been additional quorum-sensing system components identified in *V. cholerae*, including the CqsR receptor, VqmA receptor, and DPO autoinducer synthase Tdh (Fig. 1, Table 2) (Henares *et al*., 2012; Jung *et al*., 2015; Papenfort *et al*., 2017). However, while BB120 encodes homologs to some of these, their function in BB120 has not yet been tested. In order to focus specifically on similarities/differences between DS40M4 and BB120 with regard to verified functional autoinducer signaling pathways, we chose not to assay the function of these additional proteins in this manuscript. We include these as putative pathways for both BB120 and DS40M4 in Figure 1.

**Table 1.** DS40M4 homologs of established BB120 quorum-sensing proteins.

| Gene name | Amino acid identity | Query Cover | BB120 locus tag<sup>a</sup> | DS40M4 locus tag<sup>b</sup> |
| --- | --- | --- | --- | --- |
| *cqsA* | 97.96% | 100% | VIBHAR_06088 | DSB67_15960 |
| *cqsS* | 98.83% | 100% | VIBHAR_06089 | DSB67_15965 |
| *luxM* | N/A | N/A | VIBHAR_02765 | N/A |
| *luxN* | 63.23% | 99% | VIBHAR_02766 | DSB67_09895 |
| *luxP* | 97.26% | 100% | VIBHAR_05351 | DSB67_22210 |
| *luxQ* | 98.37% | 100% | VIBHAR_05352 | DSB67_22215 |
| *luxS* | 98.84% | 100% | VIBHAR_03484 | DSB67_12735 |
| *luxU* | 96.49% | 100% | VIBHAR_02958 | DSB67_10605 |
| *luxO* | 98.34% | 100% | VIBHAR_02959 | DSB67_10610 |
| *luxR* | 99.51% | 100% | VIBHAR_03459 | DSB67_12620 |
| *aphA* | 99.44% | 100% | VIBHAR_00046 | DSB67_14070 |
| *hqsK* | 98.29 | 92% | VIBHAR_01913 | DSB67_09490 |

a. Accession numbers: CP000789, CP000790, CP000791.
b. Accession numbers: CP030788, CP030789, CP030790

**Table 2.** BB120 and DS40M4 homologs of V. cholerae quorum-sensing receptor proteins.

| Gene name | BB120<sup>a</sup> |  |  | DS40M4<sup>b</sup> |  |  |
| --- | --- | --- | --- | --- | --- | --- |
|  | Amino acid identity to *V. cholerae* | Query Cover | Locus tag | Amino acid identity to *V. cholerae* | Query Cover | Locus tag |
| *vqmA* | 64.5% | 92% | VIBHAR_07094 | 64.5% | 92% | DSB67_18870 |
| *tdh* | 91.84% | 100% | VIBHAR_05001 | 91.55% | 100% | DSB67_20895 |
| *cqsR* | 54.76% | 98% | VIBHAR_01620 | 55.03% | 98% | DSB67_05165 |
| *vpsS* | 50.35% | 98% | VIBHAR_02306 | 65% | 39.37% | DSB67_09490 |
|  |  |  |  | 69% | 36.50% | DSB67_07055 |
|  |  |  |  | 70% | 39.49% | DSB67_07080 |
|  |  |  |  | 72% | 31.11% | DSB67_12445 |
|  |  |  |  | 69% | 37.06% | DSB67_12965 |
|  |  |  |  | 67% | 36.16% | LuxQ |
|  |  |  |  | 65% | 40.45% | DSB67_19025 |

a. Accession numbers: CP000789, CP000790, CP000791.
b. Accession numbers: CP030788, CP030789, CP030790.

To determine whether homologs of BB120 quorum-sensing sRNAs and proteins have similar functions in DS40M4, we generated numerous isogenic mutant strains, including Δ*cqsA* Δ*luxS*, Δ*luxO*, Δ*luxR*, *luxO* D47E (a LuxO phosphomimic allele (Freeman and Bassler, 1999a)), in addition to every combination of *qrr* deletion (single or combined) (Table S2). Our goal was to assay quorum sensing gene regulation in these DS40M4 strains by assessing bioluminescence production as a direct output of quorum sensing-driven LuxR production, which is a standard assay in the field (Jung *et al*., 2015, 2016; Kim *et al*., 2018). However, we observed that, unlike BB120, DS40M4 produces no bioluminescence at HCD (Fig. S1B). A genome search of the LuxCDABE protein sequences showed that DS40M4 only encodes a homolog of BB120 LuxB, one of the monomers of the bioluminescence protein luciferase (Meighen, 1991). DS40M4 lacks the *luxCDAE* genes, which encode the second monomer (LuxA) of the luciferase heterodimer, and the enzymes required for the production of tetradecanal, the substrate for luciferase (Fig. S1C) (Meighen, 1991). However, since most *Vibrio* species lack homologs to the *luxCDABE* genes and are incapable of producing bioluminescence, the use of a multi-copy plasmid expressing the BB120 *luxCDABE* genes under their native promoter is a standard method for assaying quorum sensing regulatory networks in heterologous *Vibrio* species, including *Vibrio cholerae* and *Vibrio vulnificus* (Lenz *et al*., 2005; Liu *et al*., 2006; Jung *et al*., 2015; Kim *et al*., 2018). We previously showed, using a P*_luxCDABE_-gfp* reporter plasmid, that wild-type DS40M4 activates the *luxCDABE* promoter from BB120, and this is dependent on DS40M4 LuxR (Simpson *et al*., 2019). We have since constructed a plasmid encoding the BB120 *luxCDABE* operon under control of its endogenous promoter (pCS38) for use in DS40M4, which enables us to monitor bioluminescence of various DS40M4 strains and assess LuxR regulation as a function of upstream quorum-sensing components.

In a wild-type DS40M4 strain containing pCS38 expressing *luxCDABE*, as the cells grow from LCD to HCD, bioluminescence is maximally produced at ~OD_600_=1.0 (Fig. 2A, 2B). The *luxO* D47E phosphomimic constitutively expresses Qrr sRNAs throughout the curve to inhibit LuxR production, resulting in very low bioluminescence, which we observe in both BB120 and DS40M4 (Fig. 2A, 2B). Conversely, we observed that a DS40M4 strain lacking all of the *qrr1-5* genes does not inhibit LuxR production and is constitutively bright, similar to BB120 (Fig. 2A, 2B). Deletion of *luxR* results in no bioluminescence, and complementation of *luxR* restores light production. Each of these phenotypes matches the analogous BB120 mutant phenotypes as published (Tu and Bassler, 2007). However, the DS40M4 Δ*luxO* strain has a distinct difference compared to BB120. In BB120, a Δ*luxO* mutant strain is constitutively bright and is similar to the Δ*qrr1-5* strain because Δ*luxO* cannot direct expression of the Qrrs (Fig. 2A). Likewise, complementation of *luxO* decreases bioluminescence at LCD compared to the Δ*luxO* strain (Fig. 2A). Curiously, in DS40M4, the Δ*luxO* strain is indistinguishable from wild-type (Fig. 2B), suggesting that in the absence of LuxO, this strain can still regulate bioluminescence production in a density-dependent manner. Complementation of *luxO* in the DS40M4 Δ*luxO* strain delayed light production, suggesting that LuxO regulation does still repress bioluminescence but is not required for density-dependent regulation of bioluminescence (Fig. 2B). One possibility to explain this result is that another functional LuxO is present in *V. campbellii*. Both BB120 and DS40M4 encode a second homolog with 49% identity to LuxO (VIBHAR_02228 and DSB67_07165, respectively). However, deletion of the DS40M4 *luxO* homolog in both the wild-type and Δ*luxO* strains did not alter the curve (Fig. S1D), indicating that this *luxO* homolog does not regulate bioluminescence. Given the epistatic relationship of the Qrrs to LuxO in control of bioluminescence, we propose that regulation of bioluminescence is occurring via an additional regulatory circuit outside the LuxO quorum-sensing pathway that also converges on the Qrrs.

**Figure 2.**
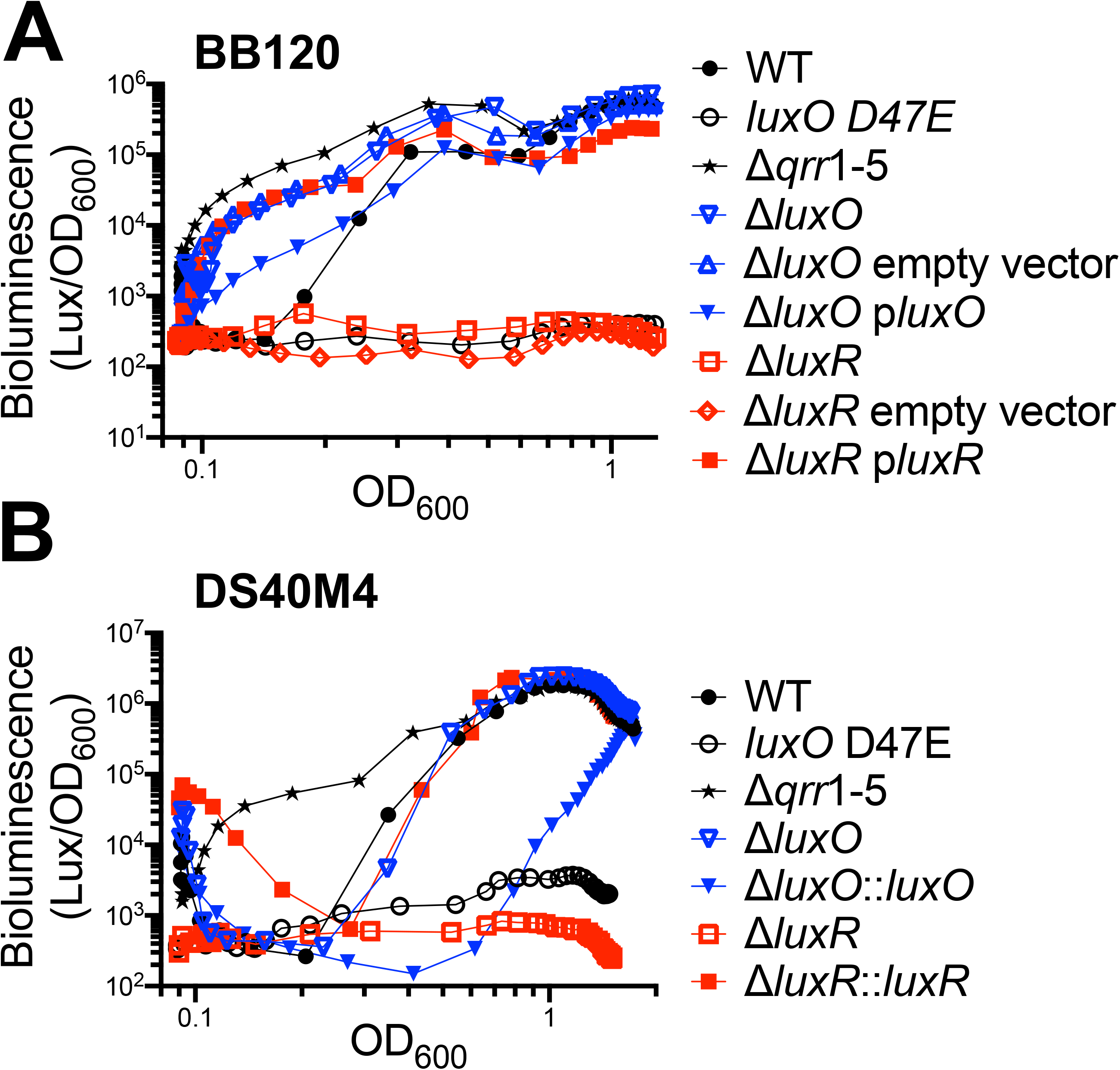
Quorum sensing regulation of bioluminescence in BB120 and DS40M4. (A, B) Bioluminescence production is shown normalized to cell density (OD_600_) during a growth curve for wild-type and mutant strains of BB120 (A) and DS40M4 containing pCS38 (B) (see Table S2 for strain names). For each graph, the data shown are from a single experiment that is representative of at least three independent biological experiments. In panel A, *luxR* and *luxO* are complemented on plasmids pKM699 and pBB147, respectively, compared to the pLAFR2 empty vector control. In panel B, *luxR* and *luxO* are complemented on the chromosome at the *luxB* locus.

### DS40M4 produces and responds to CAI-1 and AI-2 but not HAI-1

DS40M4 lacks a LuxM homolog (Fig. 3A), which is the synthase for the acyl homoserine lactone (AHL) autoinducer HAI-1 in BB120 (Bassler *et al*., 1993). However, DS40M4 encodes a LuxN homolog that is 63% identical to BB120 LuxN (Fig. 3A, 3B, Table 1, Fig. S2A). In BB120, LuxN is the HAI-1 receptor that exhibits kinase activity in the absence of HAI-1 ligand and phosphatase activity when HAI-1 is bound. Curiously, there is high sequence conservation (91% amino acid identity) between the BB120 and DS40M4 LuxN proteins in the C-terminal receiver domain (amino acids 466-849), whereas the N-terminal region of BB120 LuxN (amino acids 1-465) that binds the HAI-1 autoinducer shares only 40% identity with DS40M4 LuxN (Fig. 3B, Fig. S2A). We therefore hypothesized that DS40M4 LuxN senses a different molecule than HAI-1, possibly produced by a different synthase. DS40M4 does not encode a LuxI homolog, which is another enzyme capable of producing AHLs (Waters and Bassler, 2005). However, DS40M4 encodes close homologs of BB120 CqsS and LuxPQ (98% and 99% amino acid identity, respectively; Table 1), which bind CAI-1 and AI-2, respectively (Schauder *et al*., 2001; Ng *et al*., 2011). We investigated the activity of the autoinducers produced by DS40M4 by collecting cell-free supernatant from DS40M4 and adding it to autoinducer-sensing reporter strains in the BB120 background in which a single autoinducer synthase is deleted. For example, a Δ*cqsA* BB120 strain does not produce CAI-1 and has a lower level of bioluminescence compared to wild-type (Fig. 3C). Addition of a supernatant containing CAI-1 such as that from wild-type BB120 results in an increase in bioluminescence (Fig. 3C). We observed that addition of DS40M4 supernatant increases bioluminescence in the AI-2 and CAI-1 BB120 sensing strains (Δ*luxS* and Δ*cqsA*, respectively), but does not affect bioluminescence in the Δ*luxM* HAI-1 sensing strain (Fig. 3C). From these experiments, we conclude that DS40M4 produces AI-2 and CAI-1 molecules that can be sensed by BB120, but not an HAI-1 molecule.

**Figure 3.**
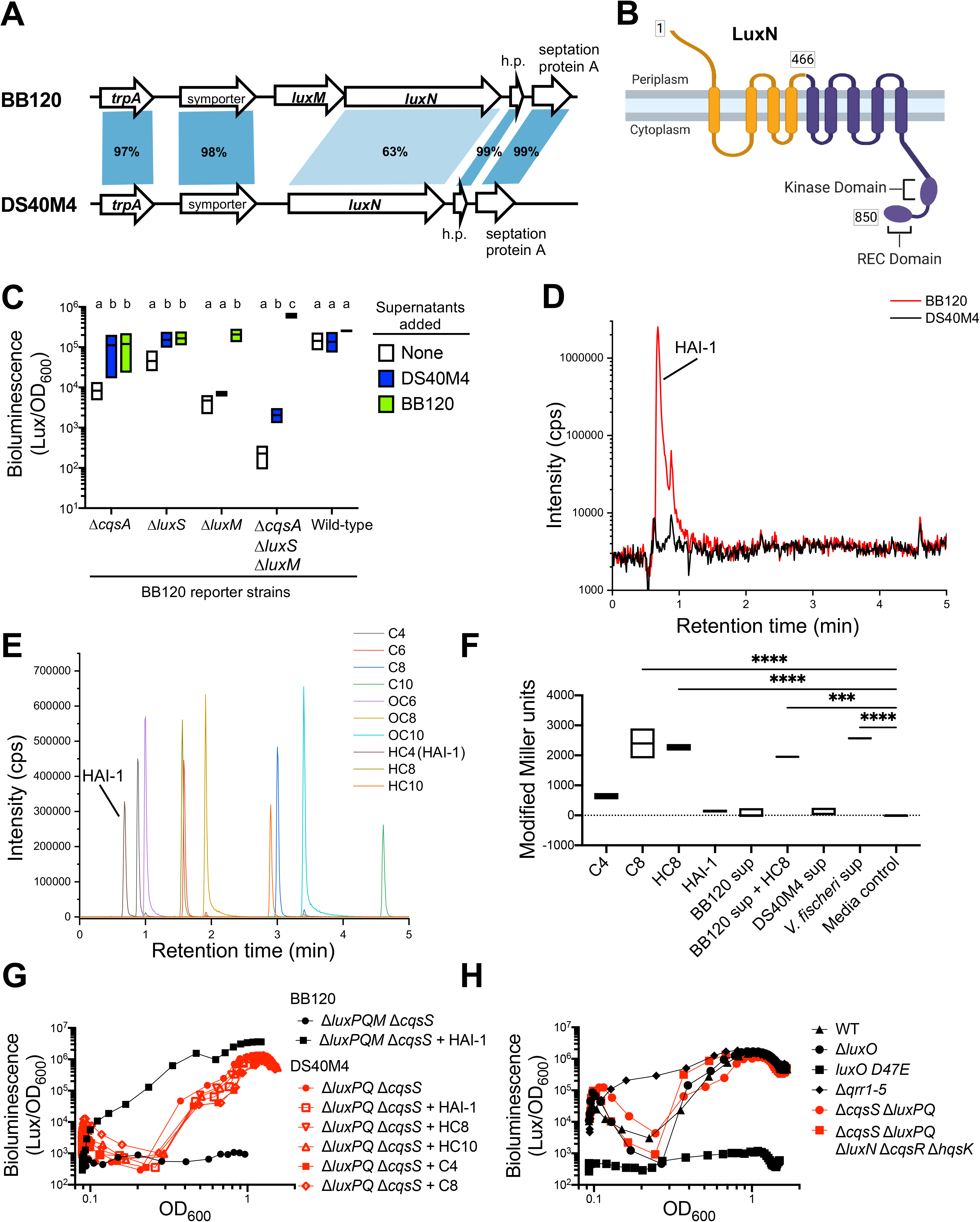
*V. campbellii* DS40M4 produces and responds to CAI-1 and AI-2 autoinducers, but not HAI-1. (A) Diagram of the *luxN* loci in BB120 and DS40M4 strains. Shaded areas indicate the percent amino acid identity shared between the strains for each gene. (B) Diagram of the predicted structure of the DS40M4 LuxN monomer. The conserved C-terminal domain is colored purple, and the N-terminal region that is not conserved is colored orange. (C) Bioluminescence production normalized to cell density (OD600) for strains in which supernatants from the indicated strains were added to BB120 strains. Different letters indicate significant differences in 2-way analysis of variance (ANOVA) of log-transformed data followed by Tukey’s multiple comparison’s test (*n* = 4, *p* < 0.05). Statistical comparisons were performed comparing the effects of the three supernatant conditions for each reporter strain; data were not compared between reporter strains. (D) Superimposed total ion chromatograms for extracts of supernatant from strains BB120 and DS40M4. (E) Superimposed extracted ion chromatograms of 10 μM AHLs. Each color corresponds to a different AHL species. (F) β-galactosidase activity as measured by Modified Miller Units in *A. tumefaciens* reporter strain KYC55 whole-cell lysate. KYC55 was grown in cell-free supernatant (sup) from *Vibrio* strains or supplemented with an exogenous AHL (10 μM) as indicated. Data shown represent at least 2 independent biological replicates except BB120 sup with spiked-in HC8 (10 μM) and *V. fischeri* sup which represent a single replicate each. (G) Bioluminescence production normalized to cell density (OD600) for BB120 Δ*luxPQM* Δ*cqsS* (TL25) and DS40M4 Δ*luxPQ* Δ*cqsS* (cas322) with or without the supplementation of exogenous AHLs (10 μM). The data shown in panels G and H are representative of three independent experiments. (H) Bioluminescence production normalized to cell density (OD600) for wild-type DS40M4 (cas291), Δ*luxO* (BDP065), *luxO* D47E (BDP062), Δ*qrr1-5* (cas269), Δ*luxPQ* Δ*cqsS* (cas322), and Δ*luxPQ* Δ*cqsS* Δ*luxN* Δ*cqsR* Δ*hqsK* (cas399).

We previously assessed the molecules sensed by DS40M4 receptors CqsS and LuxPQ (Simpson *et al*., 2019). Our data showed that DS40M4 senses the AI-2 and CAI-1 molecules produced by BB120 but does not respond to BB120 HAI-1, even though DS40M4 encodes a LuxN homolog. From both sets of data regarding autoinducers produced by BB120 and DS40M4, we conclude that DS40M4 does not produce or respond to BB120 HAI-1 under our lab conditions. To investigate whether LuxN responds to a different autoinducer, we first assayed for the presence of AHLs in DS40M4 supernatants. Using liquid chromatography - mass spectrometry (LCMS), we observed a clear signal for HAI-1 in the supernatant of BB120, which aligned as expected with the retention time of the HAI-1 peak from the synthetic control sample (Fig. 3D, 3E; hydroxy-C4-homoserine lactone (HSL); HC4). However, no signal was observed in the supernatant of DS40M4 above background intensity (Fig. 3D, S2B). To further assay for AHLs, we utilized an AHL reporter strain of *Agrobacterium tumefaciens* (strain KYC55). This reporter strain relies on the overexpression of the innate *A. tumefaciens* TraR protein that promiscuously detects AHL variants at nanomolar concentrations using β-galactosidase activity as a standard reporter (Zhu *et al*., 2003; Joelsson and Zhu, 2006). However, TraR does not respond to other classes of autoinducers, and thus β-galactosidase activity should not be induced by AI-2 or CAI-1. We observe significant activity when KYC55 is supplemented with exogenous C8 or hydroxy-C8 HSL (Fig. 3F). Surprisingly, the KYC55 reporter strain does not respond to C4 nor HAI-1, either in the supernatant of BB120 or synthetically produced (Fig. 3F). Importantly, no β-galactosidase activity is observed when the reporter strain is grown in medium supplemented with cell-free supernatants from wildtype DS40M4 relative to the LM medium control (Fig. 3F). In contrast, cell-free supernatant from *Vibrio fischeri* MJ1, which produces both C8 HSL and oxo-C6 HSL, results in a significant increase in β-galactosidase activity. Further, addition of HC8-HSL to BB120 supernatant prior to filtration also produced β-galactosidase activity, confirming that HSLs from either biological or synthetic sources can be recovered and detected from our supernatant filtration process. These data, along with conclusions drawn from the LCMS experiments, suggest that DS40M4 does not produce a detectable AHL autoinducer.

To assess whether LuxN binds to a different AHL ligand other than HAI-1, we constructed mutant strain backgrounds to specifically focus on the kinase/phosphatase activity contributed solely by LuxN. In *V. campbellii* BB120, deletion of the LuxPQ and CqsS receptors and the LuxM synthase (Δ*luxPQ* Δ*luxM* Δ*cqsS*) is sufficient to abolish quorum sensing throughout all cell densities and shut off bioluminescence (Fig. 3G). Constitutive light production can be induced in this strain when exogenous HAI-1 is supplemented (Fig. 3G). Thus, we made a Δ*luxPQ* Δ*cqsS* DS40M4 strain because we hypothesized that, in the absence of these two receptors, if LuxN is not detecting a ligand, this strain could not respond to changes in cell density. Surprisingly, the Δ*luxPQ* Δ*cqsS* strain produces bioluminescence in response to increases in cell density much like wild-type throughout the entire curve (Fig. 3G, 3H). To examine if addition of exogenous AHLs could stimulate earlier light production, we added synthetic AHLs to cultures of DS40M4 Δ*luxPQ* Δ*cqsS* at LCD and monitored bioluminescence. Neither the addition of HAI-1 nor any of the other AHL derivatives results in earlier light production for the DS40M4 Δ*luxPQ* Δ*cqsS* mutant (Fig. 3G). From these data, we conclude that DS40M4 is producing and sensing another molecule outside the LuxSPQ and CqsAS pathways. Further, if LuxN is detecting a molecule, it is not likely an AHL produced by DS40M4.

To determine what pathway(s) were functional for quorum sensing signaling, we used MuGENT to construct combinations of deletions in all genes encoding predicted quorum sensing receptors: *cqsS*, *luxPQ*, *luxN*, *cqsR*, and *hqsK*. However, the sextuple mutant strain (Δ*cqsS*, Δ*luxPQ*, Δ*luxN*, Δ*cqsR*, Δ*hqsK*) still responds to increases in cell density similar to wild266 type (Fig. 3H). These data suggest that another signal-receptor pair are functioning to drive the quorum sensing response in the absence of all predicted pathways.

### DS40M4 LuxN positively regulates bioluminescence and biofilm formation at HCD

Our collective data from our previous publication and this study show that DS40M4 does not sense or produce an AHL, but did not address whether the DS40M4 LuxN protein is a functional protein (Simpson *et al*., 2019). To determine whether DS40M4 LuxN is functional to alter regulation of gene expression downstream, we generated numerous mutants in the DS40M4 background and examined bioluminescence and biofilm formation. First, we examined the biofilm phenotype in various DS40M4 isogenic strains compared to BB120 strains using culture tubes and 96-well plates. We normalized the optical density reading after crystal violet staining (OD_590_) to the optical density for the culture prior to processing (OD_600_) because there is a large difference in growth in cultures of BB120 and DS40M4 grown statically or with shaking (Fig. 4A, S1A). In both BB120 and DS40M4, the *luxO* D47E phosphomimic allele increases biofilms above wild-type levels (Fig. 4A, 4B). Deletion of *luxN* in BB120 significantly decreases biofilms, whereas we observe that the DS40M4 Δ*luxN* strain produces the same biofilm levels as wild-type (Fig. 4A, 4B). However, based on the visual inspection of the crystal violet staining, strains BB120 Δ*luxN*, DS40M4 wild-type, and DS40M4 Δ*luxN* are nearing the lower limit of detection for this assay, thus it is possible we would not be able to detect a significant decrease in biofilm formation in DS40M4.

**Figure 4.**
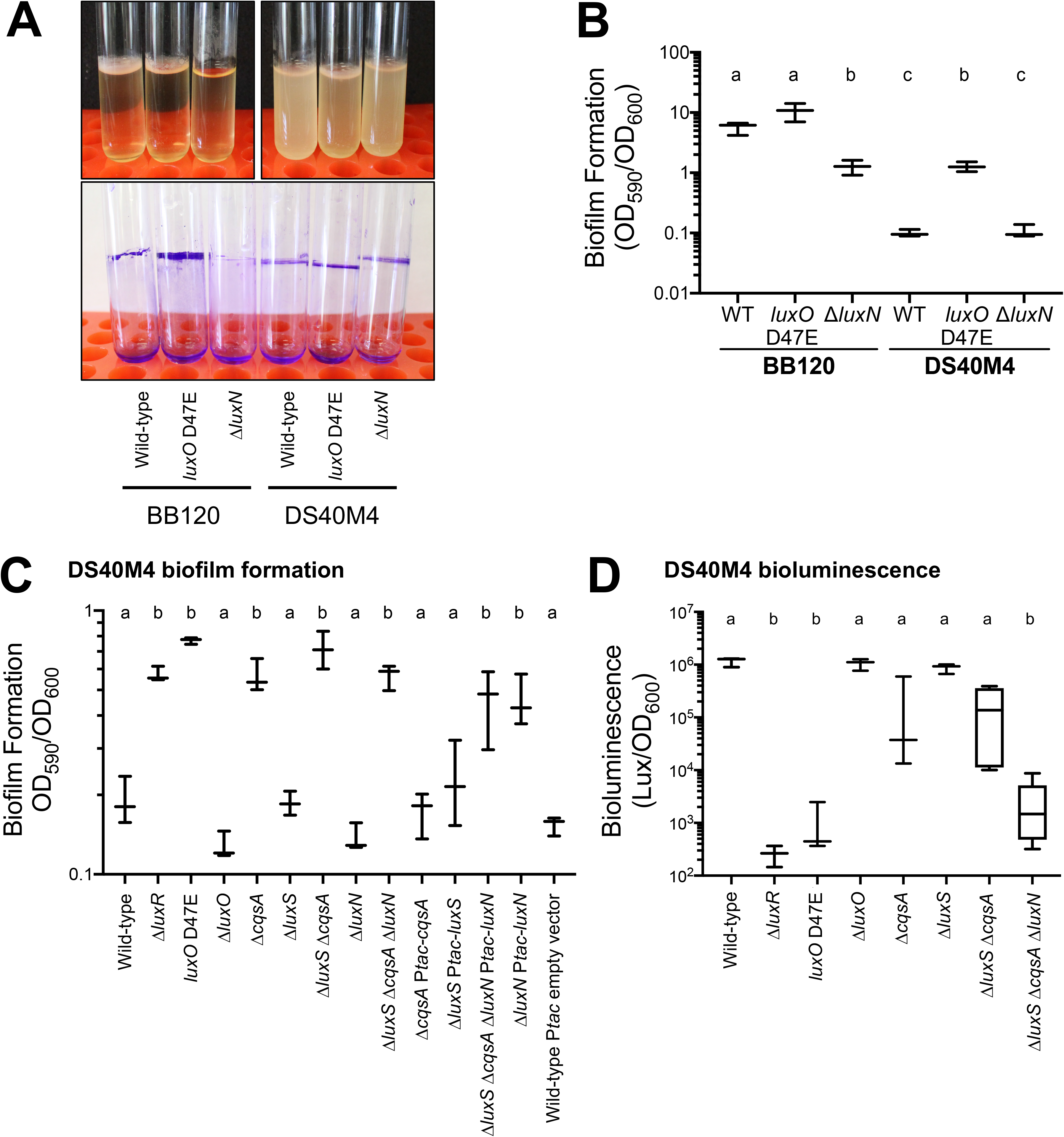
DS40M4 LuxN positively regulates biofilm formation and bioluminescence. (A) Biofilm formation measured by crystal violet staining (OD_590_) normalized to cell density (OD_600_). Culture tubes grown statically and assayed for biofilm formation by crystal violet staining. (B, C) Biofilm formation measured by crystal violet staining (OD_590_) normalized to cell density (OD_600_). Strains containing complementation plasmids or empty vector were induced with 10 μM IPTG. (D) Bioluminescence production normalized to cell density (OD_600_). For panels B, C, and D, different letters indicate significant differences in 2-way ANOVA of log-transformed data followed by Tukey’s multiple comparison’s test (*n* = 3 or *n* = 4, *p* < 0.05). See Table S2 for strain names for all panels.

To further assess the phenotypic effects of deleting *luxN* in the DS40M4 strain, we introduced the Δ*luxN* allele into a Δ*cqsA* Δ*luxS* strain. Our logic was that deletion of these two autoinducer synthases would lead to increased kinase activity of CqsS and LuxPQ, which we predicted would result in higher levels of phosphorylated LuxO and increased biofilm formation. Thus, we predicted that the effect of deleting *luxN* would be more easily observable in this background. In DS40M4, deletion of *luxR* does not maximize biofilm production to the level of the *luxO* D47E strain (Fig. 4C), possibly due to production of AphA and/or Qrrs that may still control expression of biofilm genes in a Δ*luxR* background but would not in the *luxO* D47E strain. As we predicted, the DS40M4 Δ*cqsA* Δ*luxS* mutant is significantly increased in biofilm formation compared to wild-type (Fig. 4C). Deletion of *cqsA* but not *luxS* increases biofilm formation; complementation of *cqsA* restores biofilm levels to that of wild-type DS40M4 (Fig. 4C). We observe that deletion of *luxN* in the wild-type background or the Δ*cqsA* Δ*luxS* background does not significantly alter biofilm formation, though there is a modest decrease (Fig. 4C). However, over-expression of *luxN* in the Δ*luxN* background significantly increases biofilm formation. This result, together with the observed decrease in biofilm formation in Δ*luxN* strains, indicates that LuxN is acting to positively regulate biofilm formation. However, complementation of *luxN* in the Δ*cqsA* Δ*luxS* Δ*luxN* background does not significantly alter biofilm formation (Fig. 4C), possibly because the kinase activities of CqsS and LuxPQ are dominant under the conditions in this assay.

Second, we examined the bioluminescence phenotypes of the DS40M4 isogenic deletion strains. Because the P*_tac_-luxN* complementation experiment utilized the same plasmid backbone as the bioluminescence reporter pCS38, we were unable to assay bioluminescence in the complementation strains. As shown previously (Fig. 2B), we see complete loss of bioluminescence in the DS40M4 Δ*luxR* strain. Deletion of *cqsA* or *luxS* alone does not significantly alter bioluminescence, but we note a general decrease in bioluminescence in the Δ*cqsA* strain as well as an increase in variability in the assay that we also see in the Δ*cqsA* Δ*luxS* strain (Fig. 4D). In BB120, a Δ*luxN* strain does not alter bioluminescence compared to wild-type at any point in the growth curve, but if LuxN is the only receptor expressed (Δ*luxQ* Δ*cqsS*), bioluminescence increases at LCD (Bassler *et al*., 1993; J. M. Henke and Bassler, 2004). Thus, if the DS40M4 LuxN protein functions similarly to BB120 LuxN, we predicted that deletion of *luxN* would result in decreased bioluminescence due to loss of phosphatase activity. We assayed this in the Δ*cqsA* Δ*luxS* Δ*luxN* background, which is not producing AI-2 or CAI-1, and thus the LuxPQ and CqsS receptors are predicted to be functioning as kinases at HCD. Indeed, deletion of *luxN* in the Δ*cqsA* Δ*luxS* background significantly decreases bioluminescence (Fig. 4D). From these collective data, we conclude that DS40M4 LuxN positively regulates both bioluminescence and biofilm formation. However, these experiments cannot conclusively determine whether LuxN is acting solely as a phosphatase or has kinase activity in other conditions.

### Conservation of LuxM and LuxN is widely different among vibrios

The collective results of the supernatant assays and examination of the *luxN* locus suggested that DS40M4 evolved a different quorum-sensing pathway than BB120. We were therefore curious as to the conservation of LuxMN across the *Vibrionaceae*. We used comparative genomics to identify the presence or absence of these proteins in other *Vibrio* strains. First, we used the genomes of 287 fully sequenced *Vibrio* strains from Genbank to assemble a phylogenetic tree. Strains were grouped based on a shared set of 214 complete and non-duplicated genes identified in each genome (Fig. 5A). The organization of this tree remained the same each time we randomly chose 100 genes from the set of 214 genes to reconstruct the tree, which substantiates this assembly. We observed that the species that comprise the Harveyi clade (*V. harveyi*, *V. campbellii*, *V. parahaemolyticus*, *V. natriegens*, *V. jasicida*, *V. rotiferianus*, and *V. owensii*) are grouped similarly to previous taxonomic studies (Urbanczyk *et al*., 2013). This phylogenetic tree indicates that the most closely related strains to *V. campbellii* BB120 are DS40M4 and NBRC 15631 (also known as CAIM 519, ATCC 25920) (Fig. 5B).

**Figure 5.**
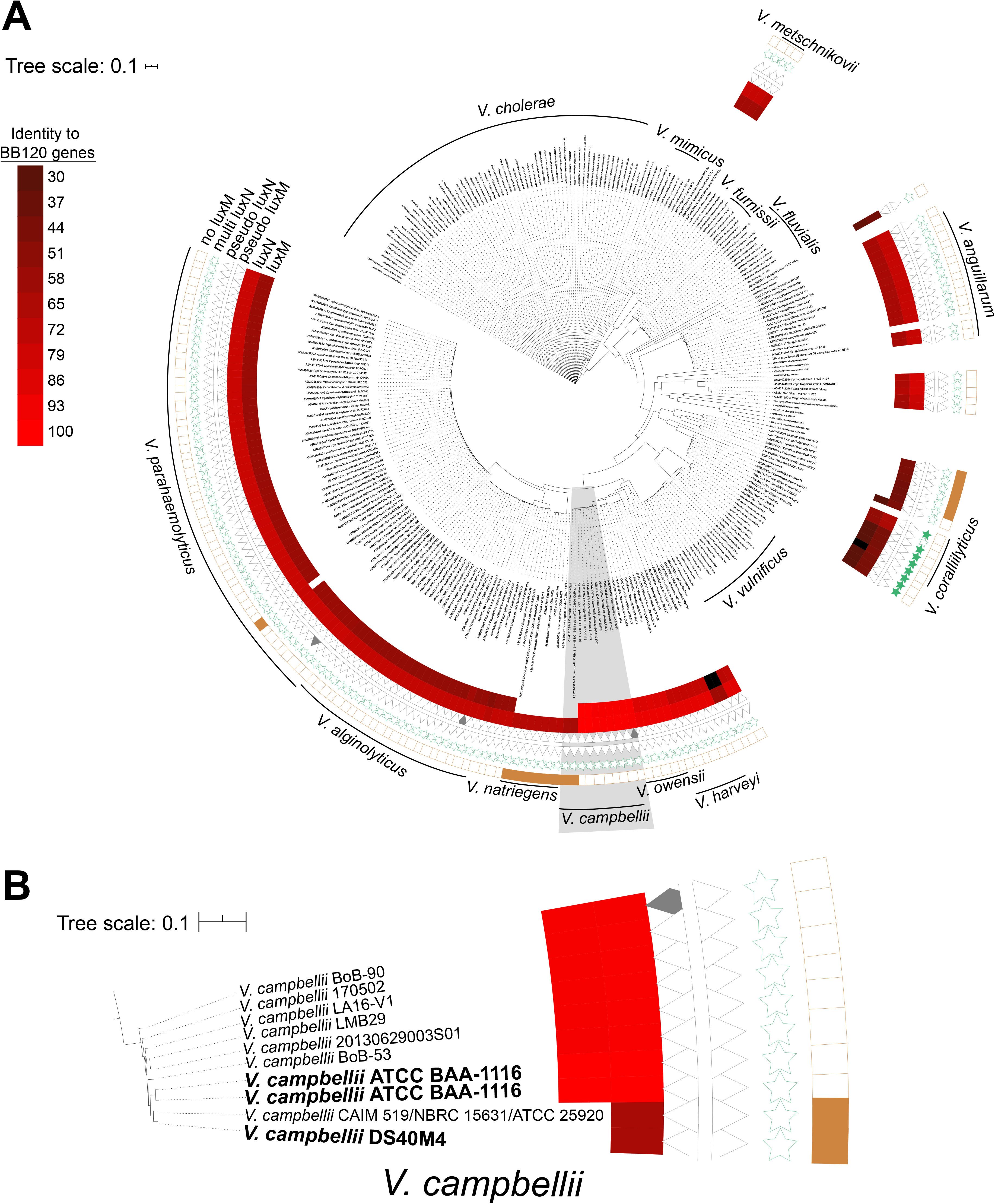
Conservation of LuxM/LuxN in *Vibrio* strains. (A) Phylogenetic tree of 287 sequenced *Vibrio* genomes from Genbank. The assembly number, genus, species, and strain are indicated for each organism. The red shading indicates the amino acid identity of a protein in these strains to either LuxM or LuxN in BB120. The presence of pseudogenes for *luxN* or *luxM*, or the presence of multiple LuxN genes is indicated by symbols. (B) Enlarged inset (from the gray region in panel A) of the *V. campbellii* region of the tree in panel A. The scale of 0.1 indicated in both panels refers to genetic distance between nodes based on nucleotide identity.

Next, we used a Hidden Markov Model to compare protein sequences from annotated and predicted genes to determine which vibrios encode either LuxM and/or LuxN (Fig. 5A). The analysis revealed that LuxMN are broadly conserved across many different *Vibrio* species and are most often found as a syntenic pair. Similar to DS40M4, our analysis showed 15 other *Vibrio* species encode LuxN but lack LuxM (2 *V. campbellii*, 1 *V. parahaemolyticus*, 7 *V. natriegens*, 2 *V. taketomensis*, 1 *V. ponticus*, 1 *V. panuliri*, and 2 *V. scophthalmi* strains). The NBRC 15631 strain also does not encode LuxM, similar to DS40M4 (Fig. 5B). In addition, several *Vibrio* strains contain LuxM or LuxN pseudogenes due to frame-shift mutations or transposon insertions, suggesting that these strains are going through reductive evolution (Fig. 5A). Conversely, some *Vibrio* strains encode multiple *luxN* genes. Together, these data show that conservation of the LuxMN system varies widely across *Vibrionaceae*. Furthermore, DS40M4 is one of only a few *Vibrio* species that encode a *luxN* homolog but not *luxM*.

### The Qrrs in DS40M4 have different activities compared to BB120

DS40M4 encodes five Qrrs with high percent nucleotide identity and similar genome position compared to BB120 (Fig. S3). Further, the Sigma-54 and LuxO binding sites identified in BB120 *qrr* promoters are conserved in DS40M4 *qrr* promoters, and only the promoter of *qrr5* diverges from BB120 at its two LuxO binding sites (Fig. S3) (Tu and Bassler, 2007). To examine the regulatory role of the Qrrs, we constructed strains containing different combinations of deletions of the *qrr1-5* genes (Table S2). We exploited the power of natural transformation in DS40M4 to construct every combination of *qrr* gene deletion, with up to four unmarked deletions in a single reaction (Fig. S4). Using MuGENT, we constructed strains expressing only a single Qrr, *e.g.*, *qrr1+*, *qrr2*+, *etc.* and assessed their bioluminescence phenotype. Surprisingly, these phenotypes also did not match previous data from BB120 (Tu and Bassler, 2007). In BB120 strains that contain only a single *qrr*, the curve exhibits increasing levels of bioluminescence depending on the Qrr gene expressed, such that the curves for each Qrr-expressing strain lie between the wild-type curve and the Δ*qrr1-5* constitutively bright curve (Fig. 6A). Specifically, in BB120, expression of Qrr2 and Qrr4 alone exhibit the strongest effect on bioluminescence repression, followed by Qrr3, then Qrr1, and finally Qrr5, which has no effect when deleted because it is not expressed under tested conditions (Fig. 6A) (Tu and Bassler, 2007). Thus, our results match those previously published in BB120: Qrr2/Qrr4>Qrr3>Qrr1>Qrr5 with regard to strength of bioluminescence.

**Figure 6.**
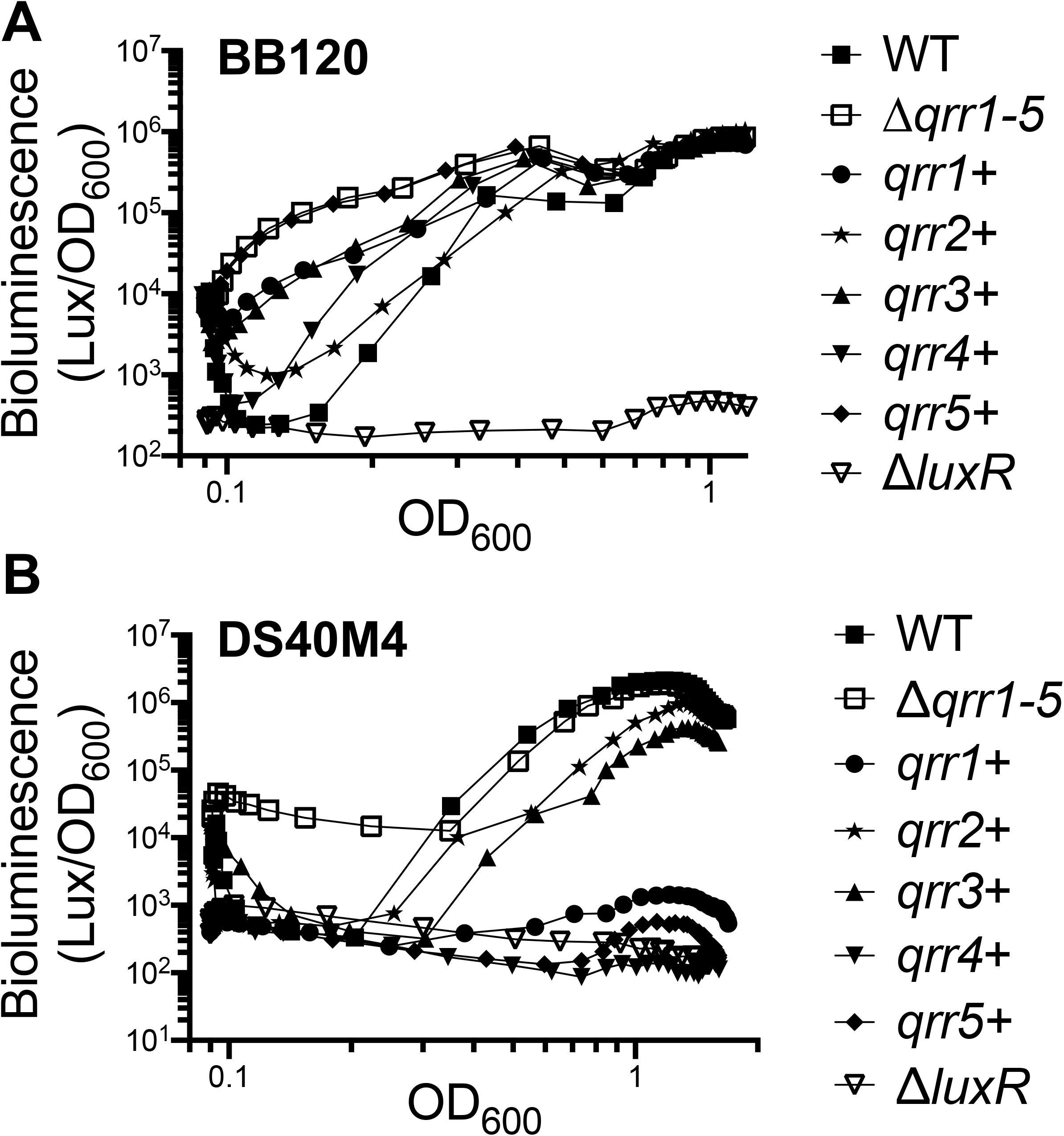
The Qrrs regulate bioluminescence in DS40M4. (A) Bioluminescence production is shown normalized to cell density (OD_600_) during a growth curve for wild-type BB120, Δ*luxR* (KM669), *qrr1*+ (KT234), *qrr2*+ (KT212), *qrr3*+ (KT225), *qrr4*+ (KT215), *qrr5*+ (KT133), and Δ*qrr1-5* (KT220). (B) Bioluminescence production is shown normalized to cell density (OD_600_) during a growth curve for wild-type DS40M4, Δ*luxR* (BP64), *qrr1*+ (cas0266), *qrr2*+ (cas0265), *qrr3*+ (cas0368), *qrr4*+ (cas0264), *qrr5*+ (cas0268), and Δ*qrr1-5* (cas0269). For both panels, a strain expressing only a single *qrr* is labeled with a ‘+’ sign. For example, *qrr4*+ expresses only *qrr4,* and the *qrr1, qrr2, qrr3,* and *qrr5* genes are deleted. The data shown for both panels are from a single experiment that is representative of at least three independent experiments.

In contrast, in DS40M4, we observe that deletion of *qrrs* has a much different effect (Fig. 6B). A strain expressing only *qrr1*, *qrr4*, or *qrr5* produces very low bioluminescence, much less than wild-type and similar to a Δ*luxR* strain (Fig. 2B, 6B). Importantly, this occurs even at HCD, when the Qrrs are not expected to be expressed based on data in BB120. Expression of only *qrr2* or *qrr3* modestly decreases bioluminescence compared to wild-type, and these strains appear to respond to the change from LCD to HCD with increasing bioluminescence production over time. (Fig. 6B). These data collectively indicate that expression of all Qrrs is not controlled only by the defined quorum-sensing pathways. However, expression of *luxO* D47E completely represses bioluminescence (Fig. 2B), suggesting that phosphorylated LuxO is epistatic to other regulatory inputs that activate bioluminescence. Further, deletion of all *qrr* genes results in constitutive bioluminescence, supporting the conclusion that the Qrrs repress bioluminescence production.

### RNA-seq defines the quorum sensing-regulated genes in DS40M4

To examine the genes regulated by quorum sensing in DS40M4, we used RNA-seq to compare gene expression of wild-type and Δ*luxR* cultures grown to OD_600_ = 1.0. We chose this optical density for several reasons: 1) this is the point at which bioluminescence is maximal and the cells have reached quorum for this phenotype in this medium (Fig. 2B), 2) this is the optical density at which RNA was collected and analyzed by RNA-seq in BB120, and 3) this density is near the end of exponential growth phase but not yet in stationary phase where stringent response might occur (Fig. S1A). Our RNA-seq analysis shows that LuxR regulates 90 genes >2-fold (*p* > 0.05, Table S1), including those involved in metabolism, cyclic di-GMP synthesis, transport, and transcriptional regulation (Table 3). This is a smaller regulon than those observed in other vibrios, including previous RNA-seq data from BB120 in which LuxR regulates >400 genes (van Kessel *et al*., 2013; Chaparian *et al*., 2016). Certain genes that are regulated by LuxR in other vibrios, such as most T6SS genes or osmotic stress genes (Wang *et al*., 2013; Shao and Bassler, 2014; Zhang *et al*., 2017; Hustmyer *et al*., 2018), are not regulated by LuxR in DS40M4. Further, the fold-change in gene activation and repression by LuxR is smaller. For example, the T3SS operons in DS40M4 are all repressed <2-fold by LuxR, whereas these are repressed >10-fold in BB120.

**Table 3.** Classes of genes regulated by LuxR in DS40M4.

| Function | Number of Genes |
| --- | --- |
| Transport | 8 |
| Metabolism | 31 |
| Virulence | 4 |
| Protease | 3 |
| Gene Expression | 4 |
| Membrane Proteins | 3 |
| Signaling | 2 |
| Oxidoreductases | 2 |
| Type VI Secretion | 1 |
| Miscellaneous | 15 |
| Hypothetical | 17 |

To extend the results of the RNA-seq analysis, we performed qRT-PCR on wild-type, Δ*luxR*, and Δ*qrr1-5* DS40M4 strains to examine the transcript levels for T3SS and T6SS, both gene classes previously shown to be regulated by LuxR in BB120 (van Kessel *et al*., 2013; Bagert *et al*., 2016; Chaparian *et al*., 2016). RNA samples were collected from these three strains at LCD (OD_600_ = 0.1), HCD (OD_600_ = 1.0), and late stationary phase (OD_600_ = 3.25), and measured by qRT-PCR. Similar to what is observed in BB120, the T3SS gene *exsB* is more highly expressed at LCD, repressed by LuxR, and activated by *qrr1-5* (Fig. 7A). In the RNA-seq data, only one T6SS gene, *tssC*, was modulated by LuxR. While qRT-PCR showed that *tssC* expression is mediated by LuxR at HCD, *tssC* is much more highly activated at late stationary phase, and this activation is not mediated by LuxR (Fig. 7B). Together, our qRT-PCR results confirmed our RNA-seq results. We conclude that the LuxR regulon in DS40M4 at HCD is smaller than that of LuxR in BB120, and the fold-change of gene regulation by LuxR is generally reduced in DS40M4.

**Figure 7.**
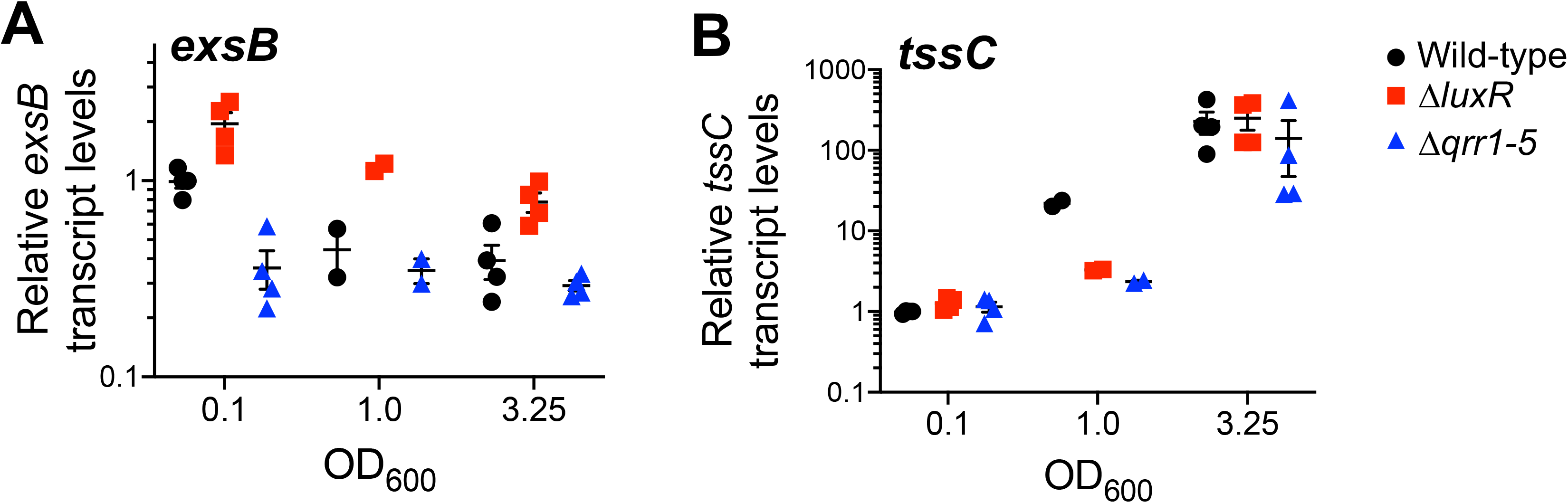
Identification of the LuxR regulon in DS40M4. (A, B) Data shown are from qRT1077 PCR analyses of transcripts of the *exsB* and *tssC* genes in the DS40M4 wild-type, Δ*luxR*, or Δ*qrr1-5* strains. Reactions were normalized to the internal standard (*hfq*).

### Virulence-associated phenotypes are differentially regulated by quorum sensing in DS40M4 and BB120

Quorum sensing plays an important role in regulating several behaviors in *Vibrio* strains that are important for *Vibrio* survival in various niches (Croxatto *et al*., 2002; Miyata *et al*., 2010; Rutherford and Bassler, 2012; Kim *et al*., 2013). To assess the impact of quorum sensing in DS40M4 and BB120, we compared three phenotypes that have previously been described as controlled by quorum sensing: interbacterial killing, exoprotease activity, and biofilm formation (Waters *et al*., 2008; Anetzberger *et al*., 2012; Shao and Bassler, 2014). BB120 exhibited stronger changes in interbacterial killing assays with quorum-sensing mutants (Fig. 8A). The LCD mutant strains, which are the *luxO* D47E strain and the Δ*luxR* strain, both showed increased killing of *E. coli* compared to wild-type. This is unexpected given previous findings in *V. campbellii* BB120 and *V. cholerae* which show that LuxR/HapR activate T6SS operons while the Qrrs repress them (Zheng *et al*., 2010; Shao and Bassler, 2014; Bagert *et al*., 2016). However, DS40M4 quorum-sensing mutants showed no significant changes in *E. coli* killing relative to the wild-type, suggesting that quorum sensing does not control this behavior in this strain (Fig. 8A). The interbacterial killing observed is likely due to the expression of T6SS genes, which is not regulated by LuxR or quorum sensing as observed in the RNA-seq and qRT-PCR data. However, it could also include production of antibacterial compounds and enzymes. Thus, we conclude that quorum sensing does not regulate interbacterial killing in DS40M4.

**Figure 8.**
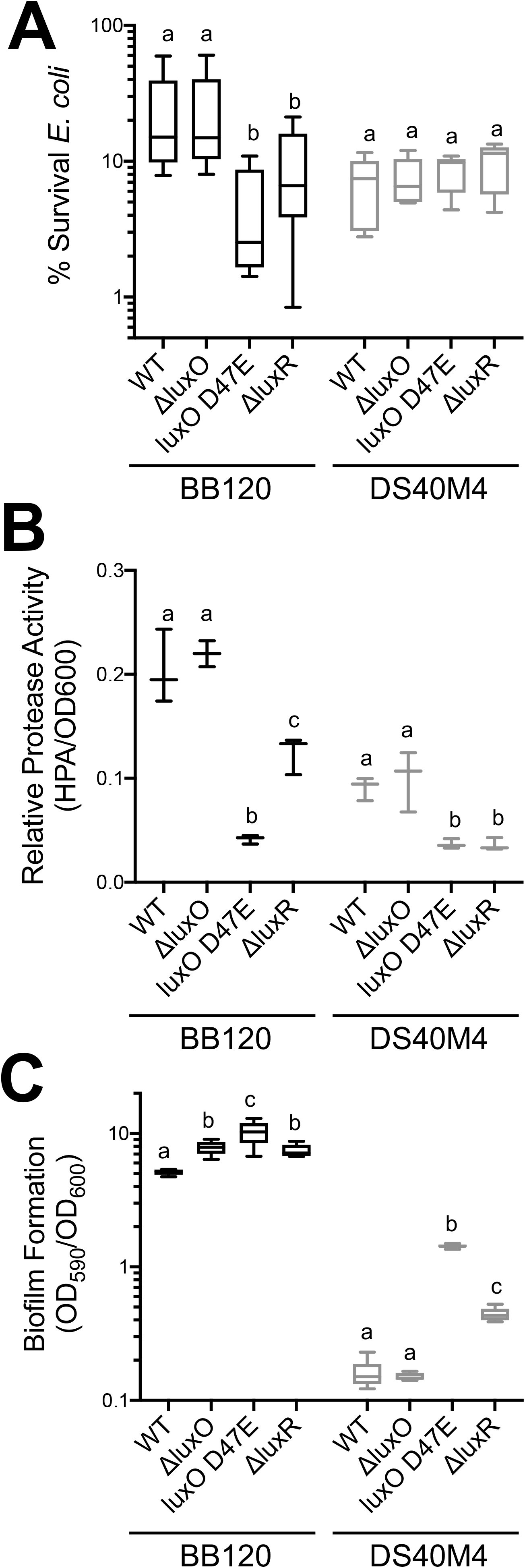
Virulence phenotypes in DS40M4 and BB120 differ in quorum-sensing regulation. For all panels, assays were performed with *V. campbellii* DS40M4 wild-type, Δ*luxO* (cas197), *luxO* D47E (BDP060), and Δ*luxR* (cas196). (A) Survival of *E. coli* in interbacterial killing assays (*n =* 9). (B) Exoprotease activity measured by HPA digestion and normalized to cell density (OD_600_). (*n* = 3). (C) Biofilm formation measured by crystal violet staining (OD_595_) normalized to cell density (OD_600_) *(n =* 6). For all panels, different letters indicate significant differences in 2-way ANOVA of log-transformed data followed by Tukey’s multiple comparison’s test (*p* < 0.05).

When we examined relative exoprotease activity between BB120 and DS40M4 strains, we observed analogous functions of the quorum-sensing proteins between the two strains. While BB120 exhibits stronger exoprotease activity than DS40M4, activity significantly drops in LCD mutants of both strains, suggesting this behavior is dependent on LuxR in both BB120 and DS40M4 (Figure 8B). Comparing biofilm formation between DS40M4 and BB120 mutants showed that LCD mutants exhibit increased biofilm formation compared to the wild-type parent strain (Figure 8C). Interestingly, however, while BB120 overall has significantly stronger biofilm formation than DS40M4, quorum sensing appears to regulate biofilm formation more strongly in DS40M4 than in BB120. From these experiments, we conclude that quorum sensing controls protease production and biofilm formation in both BB120 and DS40M4, whereas interbacterial killing is only regulated by quorum sensing in BB120.

## Discussion

*V. campbellii* BB120 has historically served as a useful model strain for quorum-sensing experiments in *Vibrio* species. BB120 was isolated as a conjugation-proficient mutant of parent strain BB7 that arose after multiple cycles of conjugation and laboratory passaging (Bonnie Bassler, personal communication). Although BB120 was originally selected as a mutant, the field has referred to BB120 as the “wild type” for ~25 years; therefore, we have kept this nomenclature (Bassler *et al*., 1997; Freeman and Bassler, 1999a, 1999b; Mok *et al*., 2003). It is perhaps not surprising that the wild isolate DS40M4 strain isolated by Haygood and colleagues (Haygood *et al*., 1993) exhibits different physiological and functional characteristics compared to BB120. DS40M4 grows at a faster rate and exhibits increased interbacterial killing of *E. coli* cells. Interestingly, however, we found that BB120 exhibits much stronger biofilm formation than DS40M4, a trait often lost by laboratory propagation (White and Surette, 2006; Davidson *et al*., 2008; McLoon *et al*., 2011), as well as a high levels of exoprotease production (Steensels *et al*., 2019). In addition, DS40M4 is one of many dark (non-bioluminescent) *V. campbellii* isolates that contain lesions in the *lux* locus that abrogate bioluminescence production (O’Grady and Wimpee, 2008). As an environmental isolate, DS40M4 likely serves as a more relevant model for comparing behaviors of *V. campbellii* to *Vibrio* strains in their native marine environment. One such behavior is the interbacterial killing activity of DS40M4. It will be informative to assess the effects of interbacterial killing and growth rate on virulence in host systems in future studies, as well as other virulence-associated characteristics such as T3SS and motility.

Although BB120 still remains a core model strain for *Vibrio* studies, it is one of several *Vibrio* strains that is not capable of natural transformation, at least not under any conditions we have tested (Simpson *et al*., 2019). Thus, making mutations in BB120 is laborious and requires construction of suicide plasmids and the use of counter-selection and colony screening. Due to this process, it can take up to three months to make a single genetic alteration. This not only drains time and resources, but disincentivizes genetic-heavy experiments as a tool for evaluating gene functions. Here we have also expanded upon our previous study of natural transformation in DS40M4 (Simpson *et al*., 2019) to show the power of natural transformation for constructing up to four unmarked mutations simultaneously (Fig. S4). The Qrr genes in the quorum-sensing circuit are only one example of a gene set for which MuGENT is useful in studying combinatorial mutant phenotypes. The ease with which mutants are constructed in DS40M4 allowed us to quickly generate deletion mutants of several core quorum-sensing genes.

Our model of the quorum-sensing circuit of DS40M4 is presented in Figure 1B based on the experiments performed in this study. DS40M4 shares the AI-2/CAI-1 pathway of BB120, but it lacks a LuxM (HAI-1 synthase) homolog, and its LuxN (HAI-1 receptor) homolog is only 63% identical. Although the DS40M4 LuxN homolog cannot sense HAI-1 from BB120, our data show that LuxN is functional and activates biofilm formation and bioluminescence. It is yet unclear if LuxN senses a ligand, even if one is produced by *V. campbellii* DS40M4. In BB120, LuxM synthesizes the AHL autoinducer HAI-1. *V. fischeri* has a LuxI synthase that produces an AHL autoinducer 3-oxo-C6; however, DS40M4 does not have a clear homolog to this protein either. Further, a protein blast against the receptor, AinR, of *V. fischeri*, which senses the autoinducer C8 AHL produced by the AinS synthase, showed 39.71% identity to the DS40M4 LuxN receptor, and no AinS homolog is present in DS40M4. Thus, it is unlikely that 3-oxo-C6 AHL or C8 AHL is the autoinducer to which DS40M4 LuxN responds. We were unable to detect any AHLs in the supernatant of DS40M4 through LCMS or *in vivo* sensor assays. Thus, we propose that DS40M4 does not produce AHLs and that LuxN activity is independent of AHL sensing. Although our experiments show that LuxN is active and regulates biofilms and bioluminescence, we could not decipher whether LuxN functions predominantly as a phosphatase or kinase using these assays. Further, the observation that BB120 Δ*luxN* exhibits low biofilm formation contradicts our prediction for this mutant based on what is seen in the literature. LuxN has been shown to be more phosphatase-biased such that when LuxN is the sole receptor expressed (Δ*luxQ* Δ*cqsS*), the strain produces more bioluminescence at LCD (J. M. Henke and Bassler, 2004). Thus, we predicted that deletion of *luxN* would yield more biofilm formation because there would be less phosphatase activity in the system, and this could yield more biofilm formation. Although we do not understand this result, it is possible that this type of assay for biofilm formation (end-point assay performed at HCD) is not appropriate for determining the function of receptors as kinases and phosphatases in which the phenotype being measured is controlled at LCD. Rather, microfluidics experiments might be a more valuable and insightful tool for this purpose as has been used for *V. cholerae* (Bridges and Bassler, 2019).

When comparing the synteny of *luxMN* across different vibrio species, we found *luxMN* are conserved as a pair in many *Vibrio* species. DS40M4 is one of a few strains, both across vibrios and more specifically within the Harveyi clade, that encodes LuxN but not LuxM. There are several evolutionary possibilities for this. One possibility is that DS40M4 never gained *luxM* compared to other members of the *V. harveyi* clade. However, another possibility is that DS40M4 lost its *luxM* gene. In our phylogeny, DS40M4 and *V. campbellii* NBRC 15631 both branch off from other members of the *V. harveyi* clade. NBRC 15631 also encodes a *luxN* but not *luxM*, and like DS40M4 its LuxN is only 63% identical to the BB120 homolog. One hypothesis suggests that DS40M4 and NBRC 15631 both lost *luxM* but maintained their *luxN* homolog. Without a cognate LuxM synthase for the LuxN receptor, it is possible that the *luxN* mutated from other *V. campbellii* homologs in such a way that it no longer responds to an autoinducer. While this hypothesis remains to be tested, future work may more clearly establish whether DS40M4 lost *luxM* from its genome.

With regard to other signaling proteins in the circuit, DS40M4 encodes an HqsK homolog with 98.29% identity to BB120 (Fig. 1B, Table 1). In BB120, HqsK has been shown to feed into the quorum-sensing pathway through interaction with a NO-sensitive H-NOX protein. Although at HCD it contributes to the dephosphorylation of LuxU, the effect of the other three receptors overwhelms the effect that NO has on bioluminescence (Henares *et al*., 2012). Since we focused this study on the effects of AI-sensing receptors on the QS pathway, we have not yet assessed the role of the HqsK homolog in DS40M4. A recently identified quorum sensing receptor protein called CqsR in *V. cholerae* has not yet been characterized in BB120. CqsR is an additional histidine sensor kinase in *V. cholerae* that phosphorylates LuxU, and thus far the ligand for this receptor has not been identified (36, 37). BB120 and DS40M4 both appear to encode homologs of *V. cholerae* CqsR with a percent identity of 54.76% and 55.03% respectively (Table 2). Another quorum-sensing system in *V. cholerae* has been identified in which the molecule DPO (3,5-dimethylpyrazin-2-ol, synthesized by threonine dehydrogenase) is bound by transcription factor VqmA (Papenfort *et al*., 2017). VqmA-DPO activates expression of a sRNA (VqmR) that represses biofilm formation. A homolog of VqmA is present in both BB120 and DS40M4 (Fig. 1, Table 2), and a putative VqmR lies directly upstream of these *vqmA* genes. Both BB120 and DS40M4 encode Tdh, which is not surprising given its role in metabolism (Table 2). However, the production of DPO, the function of VqmA and VqmR, and any connection to downstream genes still remains to be tested in most vibrios, including BB120 and DS40M4. Based on our data and the recent identification of novel autoinducers/receptors in *V. cholerae* (Jung *et al*., 2015; Papenfort *et al*., 2017), it is very possible that there are other quorum-sensing components that act in parallel or integrate into the known circuit that have yet to be identified in various *Vibrio* species. Further, the roles of HqsK, VpsS, CqsR, and LuxPQ in sensing signals from “other” cells (not exclusively self-produced) has yet to be explored in BB120 and DS40M4 (Barrasso *et al*., 2020).

Our data also suggest that DS40M4 contains a regulator that converges on the quorum-sensing circuit at the Qrrs (Fig. 1B). Contrary to BB120, deletion of *luxO* in DS40M4 does not result in maximal quorum sensing (monitored via bioluminescence production). The Δ*luxO* bioluminescence phenotype is quite similar to wild-type DS40M4, whereas the Δ*qrr*1-5 has maximal bioluminescence at all cell densities. Further complicating quorum-sensing regulation, our data also show that when either *qrr4* or *qrr5* is the only *qrr* gene remaining in DS40M4, cells have constitutively low bioluminescence levels. We propose that LuxO is not the sole regulator for the Qrrs, and that an additional regulator acts to influence Qrr expression in DS40M4. Although DS40M4 and BB120 have an additional LuxO homolog, the transcript levels of these genes are very low in both strains, and deletion of the homolog in the DS40M4 strain did not result in a constitutively bright strain. Genetic screens will likely uncover any additional regulator(s) of the Qrrs.

Previous studies of quorum sensing in *V. campbellii* have typically been performed in BB120. However, while it is understood and accepted that strains of the same bacterial species can vary widely in nature, it isn’t often investigated how the canonical models diverge amongst these strains. Although BB120 and DS40M4 are very closely related (Colston *et al*., 2019), we have shown that these two strains differ widely from one another in natural transformation, quorum-sensing pathways, and various other phenotypes. It is important to consider strain to strain variations and how bacteria in the environment may differ from the models established from laboratory strains. Here we have shown that the environmental isolate DS40M4 may serve as another useful model strain when studying behaviors in *V. campbellii*.

## Experimental Procedures

### Bacterial strains and media

All bacterial strains and plasmids are listed in Tables S2 and S3. *V. campbellii* strains were grown at 30°C on Luria marine (LM) medium (Lysogeny broth supplemented with an additional 10 g NaCl per L). Instant ocean water (IOW) medium was used in the chitin-independent transformations; it consists of Instant Ocean sea salts (Aquarium Systems, Inc.) diluted in sterile water (2X☐=☐28 g/l). Transformations were outgrown in LBv2 (Lysogeny Broth medium supplemented with additional 200☐mM NaCl, 23.14☐mM MgCl_2_, and 4.2☐mM KCl). When necessary, strains were supplemented with kanamycin (100☐μg/ml), spectinomycin (200☐μg/ml), gentamicin (100 μg/ml), or trimethoprim (10☐μg/ml). Plasmids were transferred from *E. coli* to *Vibrio* strains by conjugation on LB plates. Exconjugants were selected on LM plates with polymyxin B at 50☐U/ml and the appropriate selective antibiotic.

### Construction of linear tDNAs

All PCR products were synthesized using Phusion HF polymerase (New England Biolabs). Sequencing of constructs and strains was performed at Eurofins Scientific. Cloning procedures and related protocols are available upon request. Oligonucleotides used in the study are listed in Table S4. Linear tDNAs were generated by splicing-by-overlap extension (SOE) PCR as previously described (Ankur B. Dalia *et al*., 2014). Briefly, an UP arm containing ~3kb homology of the area upstream of the gene to be deleted was PCR amplified via a F1 and R1 primer set. Similarly, a DOWN arm containing ~3kb homology of the area downstream of the gene to be deleted was PCR amplified via a F2 and R2 primer set. For selected linear transforming DNA products, a PCR using the UP and DOWN products and an antibiotic resistance cassette as template was amplified using the F1 and R2 primers to make a selected SOE product. For unselected tDNA products, the UP and DOWN arms were used as template and amplified via the F1 and R2 primers to make a SOE product that lacked an antibiotic marker.

### Natural Transformation

DS40M4 can be induced for competence by overexpression of the master competence regulator, Tfox (chitin-independent transformation) (Simpson *et al*., 2019). Here, chitin-independent transformations were performed following the protocol established in Dalia *et al*. 2017 (Dalia *et al*., 2017). Natural competence was induced via ectopic expression of Tfox on a plasmid under an IPTG inducible promoter. A strain containing this plasmid was grown overnight in LM at 30°C, shaking, with antibiotic and 100 uM IPTG. The next day the culture was back-diluted into IOW containing 100 uM IPTG. Linear transforming DNA (tDNA) marked with an antibiotic resistance cassette was added and cells were incubated statically at 30°C for 4-6 hours. LBv2 was then added and the cells were incubated shaking at 30°C for an additional 1-2 hours. The reaction was then plated on a LM plate with the appropriate antibiotic (to select for the marker in the selected/marked tDNA) and an LM plate (to determine the viable cell count). The presence of colonies growing on the antibiotic plate containing the transformation reaction, as well as the absence of colonies growing on the antibiotic plate of the control reaction (cells induced for competence with no tDNA added) indicates a successful transformation reaction. Transformation frequency was calculated as the number of antibiotic-resistant colonies divided by viable cells. Following natural transformation, strains containing the correct target mutation were identified via colony PCR with a forward and reverse detection primer.

For MuGENT, this same process was followed, but both a marked linear tDNA and unmarked tDNA was added to the reaction. Following MuGENT, colonies were screened for integration of unselected genome edits via MASC-PCR as described previously (Ankur B Dalia *et al*., 2014). Briefly, a standard colony PCR reaction was set up, but a primer mix was added. Instead of 2 individual primers, a universal F primer was mixed with each R detection primer in equal concentrations and this mix was added to the PCR reaction. Detection primers are listed in Table S3. A band on the gel indicates the successful deletion of a targeted gene via the unselected tDNA product (Fig. S4). All mutants were further confirmed by PCR amplifying a product off of the mutant gDNA using the F1 and R2 primers, and sending the product to Eurofins Scientific for sequencing.

### Bioluminescence assays and growth curves

For the bioluminescence growth assays, overnight cultures were back-diluted 1:10,000 in 200 μl LM with selective antibiotics as needed in a black-welled clear-bottom 96-well plate (all adjacent wells left empty to avoid light carryover) and OD_600_ and bioluminescence were measured every 30 minutes for 20 hours using the BioTek Cytation 3 Plate Reader (temperature set to 30 °C, gain set to 160 for DS40M4 strains, or 135 for BB120 strains). For bioluminescence growth assays with AHLs, the AHLs were added to each well at a final concentration of 10 µM.

For endpoint bioluminescence assays, strains were back-diluted to 1:1,000 in 5 ml LM with selective antibiotics as required and incubated at 30°C shaking at 275 RPM for 7 h. 200 μl of the culture was transferred to a black-welled clear-bottom 96-well plate and the OD_600_ and bioluminescence were measured using the BioTek Cytation 3 Plate Reader (gain set to 160). For growth assays, overnight cultures were back-diluted 1:1,000 in 200 µl LM in a 96-well plate, and incubated at 30°C shaking in the BioTek Cytation 3 Plate Reader. OD_600_ was recorded every 30 minutes in the plate reader for a total of 24 hours.

For autoinducer reporter assays, overnight cultures of supernatant strains were back643 diluted 1:100 in fresh media and incubated at 30°C shaking at 275 RPM until the OD600 ~2.0. Cultures were centrifuged to pellet cells at 16,200 x g, supernatants were filtered through 0.22-μm filters, and these were added to fresh medium at 20% final concentration. Cells from strains to be assayed in the presence of supernatants were added to these supplemented media at a 1:5,000 dilution of the overnight culture and incubated shaking at 30°C at 275 RPM overnight. Bioluminescence and OD600 were measured in black-welled clear-bottom 96-well plates using the BioTek Cytation 3 Plate Reader.

### β-galactosidase reporter assays

Pre-induced cells of the ultra-sensitive *Agrobacterium tumefaciens* reporter strain KYC55 were prepared and β-galactosidase activity was measured as described previously with slight modifications (Zhu *et al*., 2003; Joelsson and Zhu, 2006). Briefly, a single aliquot of pre655 induced KYC55 cells was thawed on ice, and 1 μl/ml of cells were used to inoculate a large volume of sterile ATGN medium without antibiotics. The inoculated ATGN medium was aliquoted into 4 ml cultures containing 30% cell-free supernatant or 30% fresh LM medium supplemented with 10 μM of an exogenous AHL. Cell-free supernatants from *Vibrio* strains were prepared as described above and, as an added control, exogenous AHL was spiked into one of the cultures immediately before centrifugation to ensure AHL recovery after filtration. Cultures were incubated at 30°C at 275 RPM to an OD600 = 0.75-1.2 and the final density of each culture was recorded using a spectrophotometer. Each culture was diluted 1:40 into Z buffer and cells were lysed by the addition of 5 μl 0.05% SDS and 10 μl chloroform followed by vigorously vortexing for ~15 seconds. 200 μl of cell lysate was transferred to a clear-welled clear-bottom 96-well plate, and 20 μl of 4 mg/ml ONPG was added to each well. Immediately following the addition of ONPG, the OD420 of each well was recorded at 1-minute intervals using the BioTek Cytation 3 Plate Reader. β-galactosidase activity for each culture was calculated in Modified Miller Units as described previously (Thibodeau *et al*., 2004).

### Liquid chromatography - mass spectrometry analyses of bacterial supernatants

Filtered DS40M4 or BB120 supernatants (100 ml) were extracted with dichloromethane (3 x 30 ml). The combined organic layers were dried (MgSO4), filtered, and concentrated under reduced pressure. The dried extracts were reconstituted in 1 ml of 0.1% aqueous formic acid. Samples were injected onto an Acquity UPLC Peptide HSS T3 column (2.1 x 150 mm, 1.8 μm100 Å) by Waters with an Acquity UPLC Peptide guard column. The UPLC system consisted of an Agilent 1290 II system consisting of a multisampler, binary pump and thermostatted column compartment (set at 40°C). The components were separated by gradient elution at 0.600 ml/min. Mobile Phase A consisted of 0.1% formic acid in Optima grade water and Mobile Phase B consisted of 100% acetonitrile. All components of interest eluted within the first 5 minutes.

N-3-hydroxybutanoyl-L-homoserine lactone (HAI-1/HC4) was synthesized as previously described (Chaparian *et al*., 2019). All other AHLs were purchased from Cayman Chemical Company Inc. and resuspended in 100% acetonitrile: N-butyryl-L-homoserine lactone (C4), N683 hexanoyl-L-homoserine lactone (C6), N-octanoyl-L-homoserine lactone (C8), N-decanoyl-L684 homoserine lactone (C10), N-3-hydroxyoctanoyl-L-homoserine lactone (HC8), N-3-hydroxydecanoyl-L-homoserine lactone (HC10), N-(β-ketocaproyl)-L-homoserine lactone (OC6), N-3-oxo-octanoyl-L-homoserine lactone (OC8), and N-3-oxo-decanoyl-L-homoserine lactone (OC10). The compounds were detected qualitatively via MRM methodology on an AB Sciex QTrap 4000.

### Comparative genomics analyses

We downloaded 287 publicly available complete *Vibrio* genome sequences along with their corresponding annotated protein sequences from the Genbank database. To find any missing annotations of genes, we used Prokka version 1.12 (Seemann, 2014), a gene prediction software, by providing the annotated gene sequences as a training set. We searched the individual assemblies for *Vibrio* core genes using BUSCO version 4.02 (Seppey *et al*., 2019) against 1,445 vibrionales_odb10 HMMs. Our search returned 214 BUSCOs being marked as complete and non-duplicated in each of the 287 assemblies. A multiple sequence alignment was generated for the protein sequences matching these common core BUSCOs from each of the assemblies using MUSCLE ver. 3.8.31 (Edgar, 2004), which was then converted to nucleotide space by substituting each amino acid with the corresponding codon sequence from the associated CDS sequences. A best scoring maximum likelihood tree was drawn using RAxML version 8.2.12 (Stamatakis, 2014) with parameters (-m GTRCAT -B 0.03).

The protein sequences from annotated and predicted genes from the 287 assemblies were clustered using cd-hit ver. v4.8.1-2019-0228 (Li and Godzik, 2006) with parameters: -M 0 - g 1 -s 0.8 -c 0.65 -d 500. All sequences grouped with VIBHAR_02765 from *V. campbellii* BB120 were considered the LuxM homologs while those grouped with VIBHAR_02766 were considered LuxN homologs. To identify distant homologs with less than 65% identity, we scanned the protein sequences from the *Vibrio* assemblies against HMMs prepared from the LuxM and LuxN clusters using hmmer version 3.2.1 (http://hmmer.org/). In addition to matching scores, we also used the syntenic relationship between the hits to LuxM and LuxN HMMs to identify the corresponding homologs. The homology information was overlaid on the phylogenetic tree using the Interactive Tree of Life version 5 (Letunic and Bork, 2019).

### Extracellular protease activity assay

Extracellular protease activity was determined as previously described (Gu *et al*., 2016). Briefly, all strains were back-diluted 1:1,000 in LM medium and grown at 30°C shaking at 275 RPM for 9 h. The culture OD_600_ was measured, and the strains were collected by centrifugation at 3700 x *g* for 10 min. The supernatants of each strain were filtered through 0.22 µm filters and 1 ml of filtered supernatant was mixed with 1 ml phosphate buffer saline (PBS, pH 7.2) and 10 mg of hide powder azure (HPA). These mixtures were incubated at 37°C while shaking at 275 RPM for 2 h, and the reactions were stopped by adding 1 ml 10% trichloroacetic acid (TCA). Total protease activity was measured at 600 nm on a spectrophotometer and then normalized for each strain by dividing by the culture OD_600_.

### Biofilm formation assay

Biofilm formation was measured as previously described (O’Toole, 2010). Overnight cultures of strains grown in seawater tryptone (SWT) medium were diluted to OD_600_ = 0.005 in SWT and grown statically in culture tubes or in 96-well microtiter plates in six replicates at 30°C for 24 h. The OD_600_ of the strains in the plate were measured, and the plates were rinsed with deionized water. 0.1% crystal violet was added to each well, and the plate was subsequently rinsed with deionized water. The crystal violet-stained biofilms were then solubilized with 33% acetic acid and measured at 590 nm using the BioTek Cytation 3 Plate Reader and normalized to the OD_600_.

### Interbacterial competition assay

Interbacterial killing was measured using a competition assay as previously described (Hachani *et al*., 2013). Predator strains (*V. campbellii*) were grown in LM medium overnight at 30°C, while the prey strain (*E. coli* S17-1λpir) was grown in LB at 37°C. Predator strains were diluted 1:20 and prey strains were diluted 1:40 and grown until they reached an OD_600_ between 0.6-1.2. Cells were harvested by centrifugation at 3,700 x *g* for 10 min, and then resuspended in fresh LB to reach an OD_600_ = 10. Predator and prey strains were mixed at a ratio of 1:1, and 5 µl of each mixture was spotted onto a piece of sterile filter paper fitted to an LB plate. The competition was carried out at 30°C for 4 h, and then each filter paper was moved to 0.5 ml of LB and vortexed to resuspend the cells. These suspensions were serially diluted and plated on prey-selective plates (LB+trimethoprim_10_). Plates were incubated at 37°C until colonies were visible, then colonies were counted to determine CFU of the surviving prey.

### RNA extraction, RNA-seq, and quantitative reverse transcriptase real-time PCR (qRT-PCR)

Strains were inoculated in 5 ml LM and grown overnight shaking at 30_°_C at 275 RPM. Each strain was back-diluted 1:1,000 in LM and grown shaking at 30°C at 275 RPM until they reached an OD_600_ = 0.1, 1.0, or 3.25. Cells were collected by centrifugation at 3,700 RPM at 4°C for 10 min, the supernatant was removed, and the cell pellets were flash frozen in liquid N_2_ and stored at −80°C. RNA was isolated from pellets using a TRIzol/chloroform extraction protocol as described (Rutherford *et al*., 2011) and treated with DNase via the DNA-free^TM^ DNA Removal Kit (Invitrogen).

For RNA-seq, cDNA libraries were constructed as described previously (Papenfort *et al*., 2015) by Vertis Biotechnology AG (Freising, Germany) and sequenced using an Illumina NextSeq 500 machine in single-read mode (75 bp read length). The raw, demultiplexed reads and coverage files have been deposited in the National Center for Biotechnology Information Gene Expression Omnibus with accession code GSE147616.

qRT-PCR was used to quantify transcript levels of specific quorum sensing-controlled genes in DS40M4 and was performed using the SensiFast SYBR Hi-ROX One-Step Kit (Bioline) according to the manufacturer’s guidelines. Primers were designed to have the following parameters: amplicon size of 100 bp, primer size of 20-28 nt, and melting temperature of 55-60°C. All reactions were performed using a LightCycler 4800 II (Roche) with 0.4 µM of each primer and 5 ng of template RNA (10 µl total volume). All qRT-PCR experiments were normalized to *hfq* expression and were performed with 4 biological replicates and 2 technical replicates.

## Acknowledgements

We thank Victoria Lydick for excellent technical support, Ryan Chaparian for assistance with sequence alignments, and Konrad Förstner and Malte Siemers for support with the RNA-Seq analysis. We thank Laura Miller Conrad and Minh Tran for synthesis of HAI-1. We would also like to acknowledge the IU Arts and Sciences Undergraduate Research Experience (ASURE) for encouraging and preparing the undergraduate researchers (LG and AL) as well as Charlotte & Jim Griffin for their support of the ASURE program.

## References

Anetzberger, C., Reiger, M., Fekete, A., Schell, U., Stambrau, N., Plener, L., et al. (2012) Autoinducers Act as Biological Timers in Vibrio harveyi. PLoS One 7:.

Bagert, J.D., Van Kessel, J.C., Sweredoski, M.J., Feng, L., Hess, S., Bassler, B.L., and Tirrell, D.A. (2016) Time-resolved proteomic analysis of quorum sensing in Vibrio harveyi. Chem Sci 7: 1797–1806.

Barrasso, K., Watve, S., Simpson, C.A., Geyman, L.J., van Kessel, J.C., and Ng, W.-L. (2020) Dual-function quorum-sensing systems in bacterial pathogens and symbionts. PLOS Pathog 16: e1008934.

Bassler, B.L., Greenberg, E.P., and Stevens, A.M. (1997) Cross-species induction of luminescence in the quorum-sensing bacterium Vibrio harveyi. J Bacteriol 179: 4043 LP –4045.

Bassler, B.L., Wright, M., Showalter, R.E., and Silverman, M.R. (1993) Intercellular signalling in Vibrio harveyi: sequence and function of genes regulating expression of luminescence. Mol Microbiol 9: 773–786.

Bassler, B.L., Wright, M., and Silverman, M.R. (1994) Sequence and function of LuxO, a negative regulator of luminescence in Vibrio harveyi. Mol Microbiol 12: 403–412.

Bridges, A.A. and Bassler, B.L. (2019) The intragenus and interspecies quorum-sensing autoinducers exert distinct control over Vibrio cholerae biofilm formation and dispersal. PLoS Biol 17: 1–28.

Chaparian, R.R., Olney, S.G., Hustmyer, C.M., Rowe-Magnus, D.A., and van Kessel, J.C. (2016) Integration host factor and LuxR synergistically bind DNA to coactivate quorum-sensing genes in Vibrio harveyi. Mol Microbiol 101: 823–840.

Chaparian, R.R., Tran, M.L.N., Miller Conrad, L.C., Rusch, D.B., and van Kessel, J.C. (2019) Global H-NS counter-silencing by LuxR activates quorum sensing gene expression. Nucleic Acids Res 1–13.

Colston, S.M., Hervey, W.J., Horne, W.C., Haygood, M.G., Petersen, B.D., van Kessel, J.C., and Vora, G.J. (2019) Complete Genome Sequence of Vibrio campbellii DS40M4. Microbiol Resour Announc 8: 1–2.

Croxatto, A., Chalker, V.J., Lauritz, J., Jass, J., Hardman, A., Williams, P., et al. (2002) VanT, a homologue of Vibrio harveyi LuxR, regulates serine, metalloprotease, pigment, and biofilm production in Vibrio anguillarum. J Bacteriol 184: 1617–1629.

Dalia, A.B. (2018) Natural Cotransformation and Multiplex Genome Editing by Natural Transformation (MuGENT) of Vibrio cholerae. In Vibrio Cholerae: Methods and Protocols. Sikora, A.E. (ed). New York, NY: Springer New York, pp. 53–64.

Dalia, Ankur B., Lazinski, D.W., and Camilli, A. (2014) Identification of a membrane-bound transcriptional regulator that links chitin and natural competence in Vibrio cholerae. MBio 5:.

Dalia, Ankur B, McDonough, E., and Camilli, A. (2014) Multiplex genome editing by natural transformation. Proc Natl Acad Sci 111: 8937–8942.

Dalia, T.N., Hayes, C.A., Stolyar, S., Marx, C.J., McKinlay, J.B., and Dalia, A.B. (2017) Multiplex Genome Editing by Natural Transformation (MuGENT) for Synthetic Biology in Vibrio natriegens. ACS Synth Biol 6: 1650–1655.

Davidson, C.J., White, A.P., and Surette, M.G. (2008) Evolutionary loss of the rdar morphotype in Salmonella as a result of high mutation rates during laboratory passage. ISME J 2: 293– 307.

Edgar, R.C. (2004) MUSCLE: A multiple sequence alignment method with reduced time and space complexity. BMC Bioinformatics 5: 1–19.

Freeman, J.A. and Bassler, B.L. (1999a) A genetic analysis of the function of LuxO, a two-component response regulator involved in quorum sensing in Vibrio harveyi. Mol Microbiol 31: 665–677.

Freeman, J.A. and Bassler, B.L. (1999b) Sequence and function of LuxU: A two-component phosphorelay protein that regulates quorum sensing in Vibrio harveyi. J Bacteriol 181: 899–906.

Freeman, J.A., Lilley, B.N., and Bassler, B.L. (2000) A genetic analysis of the functions of LuxN: A two-component hybrid sensor kinase that regulates quorum sensing in Vibrio harveyi. Mol Microbiol 35: 139–149.

Gode-Potratz, C.J. and McCarter, L.L. (2011) Quorum sensing and silencing in Vibrio parahaemolyticus. J Bacteriol 193: 4224–4237.

Gu, D., Guo, M., Yang, M., Zhang, Y., Zhou, X., and Wang, Q. (2016) A σ Temperature Gauge Controls a Switch from LuxR-Mediated Virulence Gene Expression to Thermal Stress Adaptation in Vibrio alginolyticus. PLoS Pathog 12: 1–31.

Hachani, A., Lossi, N.S., and Filloux, A. (2013) A visual assay to monitor T6SS-mediated bacterial competition. J Vis Exp e50103.

Haygood, M.G., Holt, P.D., and Butler, A. (1993) Aerobactin production by a planktonic marine Vibrio sp. Limnol Oceanogr 38: 1091–1097.

Henares, B.M., Higgins, K.E., and Boon, E.M. (2012) Discovery of a nitric oxide responsive quorum sensing circuit in vibrio harveyi. ACS Chem Biol 7: 1331–1336.

Henke, J. and Bassler, B. (2004) Quorum sensing regulates type III secretion in Vibrio harveyi and Vibrio parahaemolyticus. J Bacteriol 186: 3794–3805.

Henke, J.M. and Bassler, B.L. (2004) Three Parallel Quorum-Sensing Systems Regulate Gene Expression in *Vibrio harveyi* J Bacteriol 186: 6902 LP – 6914.

Hustmyer, C.M., Simpson, C.A., Olney, S.G., Rusch, D.B., Bochman, M.L., and van Kessel, J.C. (2018) Promoter Boundaries for the luxCDABE and betIBA-proXWV Operons in Vibrio harveyi Defined by the Method Rapid Arbitrary PCR Insertion Libraries (RAIL). J Bacteriol 200: e00724–17.

Joelsson, A.C. and Zhu, J. (2006) LacZ-Based Detection of Acyl-Homoserine Lactone Quorum-Sensing Signals. Curr Protoc Microbiol 3: 1C.2.1–1C.2.9.

Jung, S.A., Chapman, C.A., and Ng, W.L. (2015) Quadruple Quorum-Sensing Inputs Control Vibrio cholerae Virulence and Maintain System Robustness. PLoS Pathog 11: 1–19.

Jung, S.A., Hawver, L.A., and Ng, W.L. (2016) Parallel quorum sensing signaling pathways in Vibrio cholerae. Curr Genet 62: 255–260.

van Kessel, J.C., Rutherford, S.T., Shao, Y., Utria, A.F., and Bassler, B.L. (2013) Individual and combined roles of the master regulators apha and luxr in control of the Vibrio harveyi quorum-sensing regulon. J Bacteriol 195: 436–443.

Kim, B.S., Jang, S.Y., Bang, Y., Hwang, J., Koo, Y., Jang, K.K., et al. (2018) QStatin, a Selective Inhibitor of Quorum Sensing in Vibrio Species. MBio 9: e02262–17.

Kim, S.M., Park, J.H., Lee, H.S., Kim, W. Bin, Ryu, J.M., Han, H.J., and Choi, S.H. (2013) LuxR Homologue SmcR Is Essential for Vibrio vulnificus Pathogenesis and Biofilm Detachment, and Its Expression is Induced by Host Cells. Infect Immun 81: 3721 LP –3730.

Lenz, D.H., Miller, M.B., Zhu, J., Kulkarnl, R. V., and Bassler, B.L. (2005) CsrA and three redundant small RNAs regulate quorum sensing in Vibrio cholerae. Mol Microbiol 58: 1186–1202.

Letunic, I. and Bork, P. (2019) Interactive Tree of Life (iTOL) v4: Recent updates and new developments. Nucleic Acids Res 47: 256–259.

Li, W. and Godzik, A. (2006) Cd-hit: A fast program for clustering and comparing large sets of protein or nucleotide sequences. Bioinformatics 22: 1658–1659.

Lin, B., Wang, Z., Malanoski, A.P., O’Grady, E.A., Wimpee, C.F., Vuddhakul, V., et al. (2010) Comparative genomic analyses identify the Vibrio harveyi genome sequenced strains BAA-1116 and HY01 as Vibrio campbellii. Environ Microbiol Rep 2: 81–89.

Liu, Z., Hsiao, A., Joelsson, A., and Zhu, J. (2006) The transcriptional regulator VqmA increases expression of the quorum-sensing activator HapR in Vibrio cholerae. J Bacteriol 188: 2446–2453.

McLoon, A.L., Guttenplan, S.B., Kearns, D.B., Kolter, R., and Losick, R. (2011) Tracing the domestication of a biofilm-forming bacterium. J Bacteriol 193: 2027–2034.

Meibom, K.L., Blokesch, M., Dolganov, N.A., Wu, C.Y., and Schoolnik, G.K. (2005) Microbiology: Chitin induces natural competence in vibrio cholerae. Science (80-) 310: 1824–1827.

Meighen, E.A. (1991) Molecular biology of bacterial bioluminescence. Microbiol Rev 55: 123– 142.

Miyata, S.T., Kitaoka, M., Wieteska, L., Frech, C., Chen, N., and Pukatzki, S. (2010) The Vibrio Cholerae Type VI Secretion System: Evaluating its Role in the Human Disease Cholera. Front Microbiol 1: 117.

Mok, K.C., Wingreen, N.S., and Bassler, B.L. (2003) Vibrio harveyi quorum sensing: A coincidence detector for two autoinducers controls gene expression. EMBO J 22: 870–881.

Ng, W.L., Perez, L.J., Wei, Y., Kraml, C., Semmelhack, M.F., and Bassler, B.L. (2011) Signal production and detection specificity in Vibrio CqsA/CqsS quorum-sensing systems. Mol Microbiol 79: 1407–1417.

O’Grady, E.A. and Wimpee, C.F. (2008) Mutations in the lux operon of natural dark mutants in the genus vibrio. Appl Environ Microbiol 74: 61–66.

O’Toole, G.A. (2010) Microtiter dish Biofilm formation assay. J Vis Exp 10–11.

Papenfort, K., Förstner, K.U., Cong, J.P., Sharma, C.M., and Bassler, B.L. (2015) Differential RNA-seq of Vibrio cholerae identifies the VqmR small RNA as a regulator of biofilm formation. Proc Natl Acad Sci U S A 112: E766–E775.

Papenfort, K., Silpe, J.E., Schramma, K.R., Cong, J.-P., Seyedsayamdost, M.R., and Bassler, B.L. (2017) A Vibrio cholerae autoinducer–receptor pair that controls biofilm formation. Nat Chem Biol 13: 551–557.

Rutherford, S.T. and Bassler, B.L. (2012) Bacterial quorum sensing: Its role in virulence and possibilities for its control. Cold Spring Harb Perspect Med 2: 1–25.

Rutherford, S.T., van Kessel, J.C., Shao, Y., and Bassler, B.L. (2011) AphA and LuxR/HapR reciprocally control quorum sensing in vibrios. Genes Dev 25: 397–408.

Schauder, S., Shokat, K., Surette, M.G., and Bassler, B.L. (2001) The LuxS family of bacterial autoinducers: Biosynthesis of a novel quorum-sensing signal molecule. Mol Microbiol 41: 463–476.

Seemann, T. (2014) Prokka: Rapid prokaryotic genome annotation. Bioinformatics 30: 2068– 2069.

Seppey, M., Manni, M., and Zdobnov, E.M. (2019) BUSCO: Assessing Genome Assembly and Annotation Completeness. Methods Mol Biol 1962: 227–245.

Shao, Y. and Bassler, B.L. (2014) Quorum regulatory small RNAs repress type VI secretion in Vibrio cholerae. Mol Microbiol 92: 921–930.

Simpson, C.A., Podicheti, R., Rusch, D.B., Dalia, A.B., and van Kessel, J.C. (2019) Diversity in Natural Transformation Frequencies and Regulation across Vibrio Species. MBio 10: 1–16.

Stamatakis, A. (2014) RAxML version 8: A tool for phylogenetic analysis and post-analysis of large phylogenies. Bioinformatics 30: 1312–1313.

Steensels, J., Gallone, B., Voordeckers, K., and Verstrepen, K.J. (2019) Domestication of Industrial Microbes. Curr Biol 29: R381–R393.

Sun, Y., Bernardy, E.E., Hammer, B.K., and Miyashiro, T. (2013) Competence and natural transformation in vibrios. Mol Microbiol 89: 583–595.

Thibodeau, S.A., Fang, R., and Joung, J.K. (2004) High-throughput β bacterial cell-based reporter systems. Biotechniques 36: 410–415.

Tu, K.C. and Bassler, B.L. (2007) Multiple small RNAs act additively to integrate sensory information and control quorum sensing in Vibrio harveyi. Genes Dev 21: 221–233.

Urbanczyk, H., Ogura, Y., and Hayashi, T. (2013) Taxonomic revision of Harveyi clade bacteria (family Vibrionaceae) based on analysis of whole genome sequences. Int J Syst Evol Microbiol 63: 2742–2751.

Wang, L., Zhou, D., Mao, P., Zhang, Y., Hou, J., Hu, Y., et al. (2013) Cell density- and quorum sensing-dependent expression of type VI secretion system 2 in vibrio parahaemolyticus. PLoS One 8: 1–11.

Waters, C.M. and Bassler, B.L. (2005) QUORUM SENSING: Cell-to-Cell Communication in Bacteria. Annu Rev Cell Dev Biol 21: 319–346.

Waters, C.M., Lu, W., Rabinowitz, J.D., and Bassler, B.L. (2008) Quorum sensing controls biofilm formation in Vibrio cholerae through modulation of cyclic Di-GMP levels and repression of vpsT. J Bacteriol 190: 2527–2536.

White, A.P. and Surette, M.G. (2006) Comparative genetics of the rdar morphotype in Salmonella. J Bacteriol 188: 8395–8406.

Yamamoto, S., Izumiya, H., Mitobe, J., Morita, M., Arakawa, E., Ohnishi, M., and Watanabe, H. (2011) Identification of a chitin-induced small RNA that regulates translation of the tfoX gene, encoding a positive regulator of natural competence in vibrio cholerae. J Bacteriol 193: 1953–1965.

Zhang, Yiquan, Gao, H., Osei-Adjei, G., Zhang, Ying, Yang, W., Yang, H., et al. (2017) Transcriptional regulation of the type VI secretion system 1 genes by quorum sensing and ToxR in Vibrio parahaemolyticus. Front Microbiol 8:.

Zheng, J., Shin, O.S., Cameron, D.E., and Mekalanos, J.J. (2010) Quorum sensing and a global regulator TsrA control expression of type VI secretion and virulence in Vibrio cholerae. Proc Natl Acad Sci 107: 21128 LP–21133.

Zhu, J., Chai, Y., Zhong, Z., Li, S., and Winans, S.C. (2003) Agrobacterium Bioassay Strain for Ultrasensitive Detection of N-Acylhomoserine Lactone-Type Quorum-Sensing Molecules: Detection of Autoinducers in Mesorhizobium huakuii. Appl Environ Microbiol 69: 6949– 6953.

