## Supplementary Info for "The quorum-sensing systems of *Vibrio campbellii* DS40M4 and BB120 are genetically and functionally distinct"

Supplemental Figures S1-S4
Supplemental Tables S1-S4

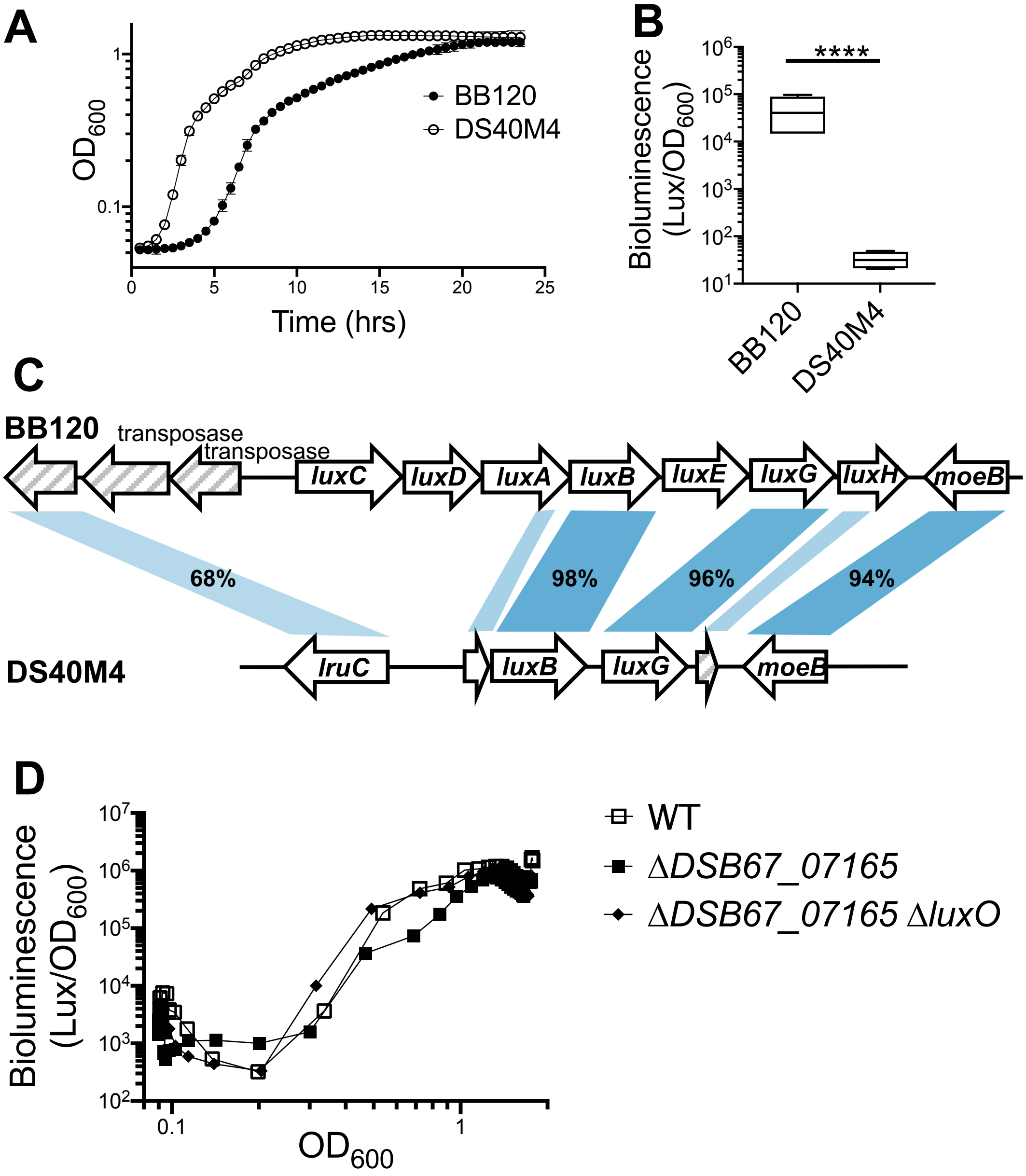

**Figure S1: Comparison of growth and bioluminescence between BB120 and DS40M4**. (A) Growth curve comparing growth rates between BB120 and DS40M4 in LM media. (B) Bioluminescence production normalized to cell density ($OD_{600}$) ($n$ = 4). (C) Diagram indicating the presence of the genes encoding LuxCDABE in BB120 and in DS40M4. For panel B, unpaired, two-tailed t-tests were performed on log-transformed data. (D) Bioluminescence production is shown normalized to cell density ($OD_{600}$) during a growth curve for wild-type DS40M4, ΔDSB67_07165 (gene encoding LuxO homolog; cas303), and ΔDSB67_07165 Δ*luxO* (cas304).

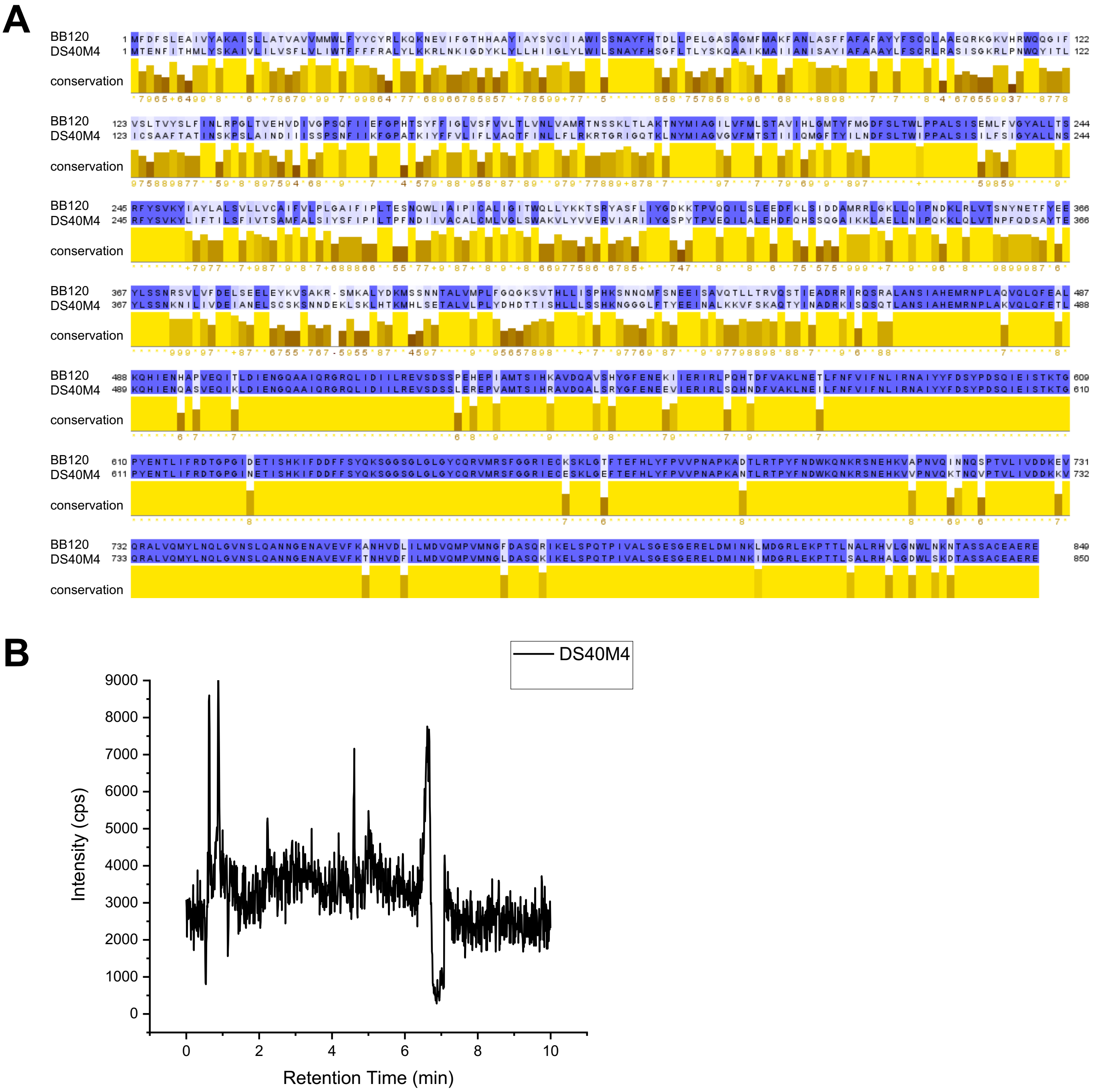

**Figure S2**. (A) Alignment of LuxN proteins from BB120 and DS40M4. Amino acid conservation is indicated by blue shading and by the yellow bars underneath. (B) Total ion chromatogram for supernatant extract from strain DS40M4.

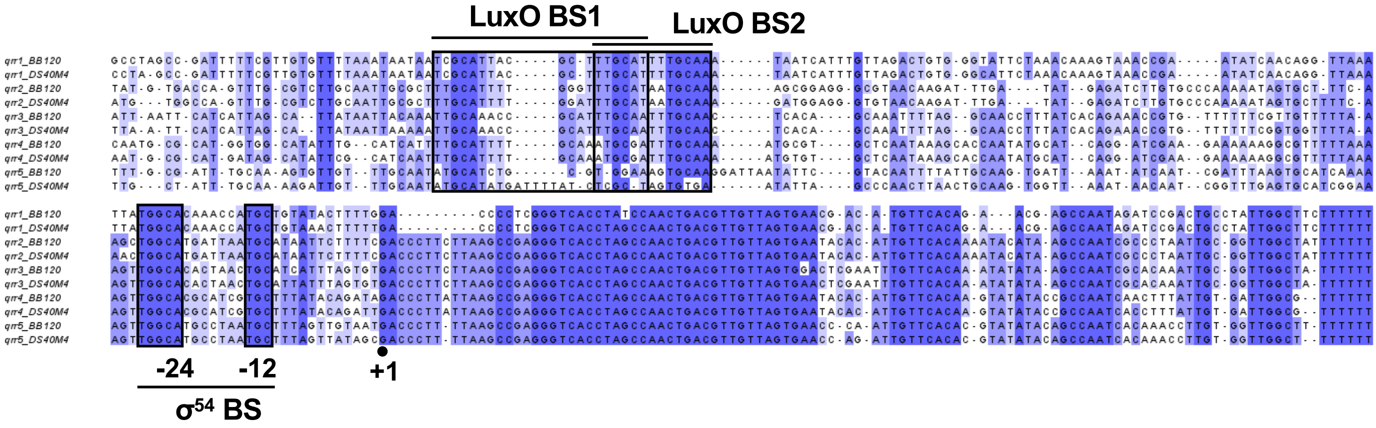

**Figure S3.** Alignment of the Qrr genes and promoters from BB120 and DS40M4. Nucleotide conservation is indicated by blue shading. Conserved binding sequences are indicated in boxes.

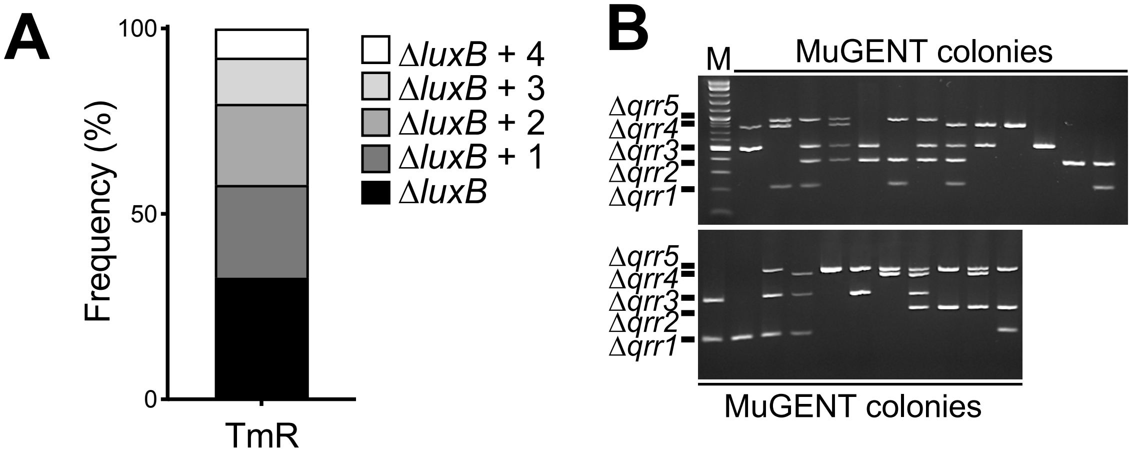

**Figure S4**. **MuGENT enables deletion of Qrr sRNAs in combinations of up to four unselected mutations.** (A) Frequency of recovery of the selectable *luxB* deletion alone or in combination with one, two, three, or four *qrr* deletions using MuGENT in DS40M4. (B) Representative agarose gel of 24 MASC-PCR products from transformations targeting all five *qrr* genes *(qrr1-5)* and *luxB* in DS40M4. The presence of a band indicates that the unmarked gene has been deleted in that colony.

**Table S1.** (See excel file.)

**Table S2. Strains used in this study.**

| Name | Description | Reference |
|---|---|---|
| BB120 | *V. campbellii* type strain, derivative of BB7 (wild-type) | (Bassler *et al.*, 1997) |
| DS40M4 | Wild-type *V. campbellii* | (Dias *et al.*, 2012) |
| cas0034 | DS40M4, pMMB67EH-tfox-kanR | (Simpson *et al.*, 2019) |
| cas0107 | DS40M4, pMMB67EH-tfox-kanR, Δl*uxB*::*spec*$^{R}$ | This study |
| cas0291 | DS40M4, Δ*luxB*::*spec*$^{R}$ , pCS38 | This study |
| cas0283 | DS40M4, pMMB67EH-tfox-kanR, Δ*luxB*::*spec*$^{R}$, Δ*cqsA*, Δ*luxS*, Δ*luxN* | This study |
| cas0252 | DS40M4, pMMB67EH-tfox-kanR, Δ*luxB*::*tm*$^{R}$, Δ*cqsA*, Δ*luxS*, | This study |
| cas0300 | DS40M4, pMMB67EH-tfox-kanR, Δ*luxB*::*spec*$^{R}$, Δ*luxN* | This study |
| cas0299 | DS40M4, Δ*luxB*::*tm*$^{R}$, *qrr4*+, Δ*luxO*::*spec*$^{R}$, + pCS38 | This study |
| cas0269 | DS40M4, Δ*luxB*::*spec*$^{R}$, *Δqrr1-5*, + pCS38 | This study |
| BDP060 | DS40M4, pMMB67EH-tfox-kanR, Δ*luxB*::*spec*$^{R}$, *luxO* D47E | This study |
| BDP062 | DS40M4, Δ*luxB*::*spec*$^{R}$, *luxO* D47E, + pCS38 | This study |
| BDP064 | DS40M4, Δ*luxB*::*spec*$^{R}$, Δ*luxR*, + pCS38 | This study |
| BDP065 | DS40M4, Δ*luxB*::*spec*$^{R}$, Δ*luxO*, + pCS38 | This study |
| BDP090 | DS40M4, pMMB67EH-tfox-kanR, Δ*luxB*::*spec*$^{R}$, Δ*cqsA* | This study |
| cas0285 | DS40M4, Δ*luxB*::*spec*$^{R}$, Δ*cqsA*, Δ*luxS*, Δ*luxN*, + pCS38 | This study |
| LG002 | DS40M4, Δ*luxB*::*spec*$^{R}$, Δ*cqsA*, Δ*luxS*, + pCS38 | This study |
| LG001 | DS40M4, Δ*luxB*::*tm*$^{R}$, Δ*luxS*, + pCS38 | This study |
| LG003 | DS40M4, Δ*luxB*::*tm*$^{R}$ , Δ*cqsA*, + pCS38 | This study |
| TL189 | BB120, Δ*luxM* Δ*cqsA* Δ*luxS* | This study |
| KM669 | BB120, Δ*luxR* | (Pompeani *et al.*, 2008) |
| cas0368 | DS40M4, Δ*luxB*::*spec*$^{R}$, *qrr3*+, + pCS38 | This study |
| cas0268 | DS40M4, Δ*luxB*::*tm*$^{R}$, *qrr5*+, + pCS38 | This study |
| cas0264 | DS40M4, Δ*luxB*::*tm*$^{R}$, *qrr4*+, + pCS38 | This study |
| cas0265 | DS40M4, Δ*luxB*::*tm*$^{R}$, *qrr2*+, + pCS38 | This study |
| cas0266 | DS40M4, Δ*luxB*::*tm*$^{R}$, *qrr1*+, + pCS38 | This study |
| BB170 | BB120*, luxN::Tn5* | (Bassler *et al.*, 1993) |
| KM816 | BB120, Δ*luxS* | (Waters and |

| | | |
|---|---|---|
| | | Bassler, 2006) |
| TL203 | BB120, Δ*cqsA* | (Simpson *et al.*, 2019) |
| JMH363 | BB120, Δ*luxM* | (Waters and Bassler, 2006) |
| JAF548 | BB120, *luxO* D47E | (Henke and Bassler, 2004) |
| S17-1λ*pir* | *E. coli*, λ-*pir*, *recA thi pro hsdR*$^{-}$ *M*$^{+}$ RP4: 2-Tc, Mu, Tp$^{R}$, Sm$^{R}$ | (de Lorenzo and Timmis, 1994) |
| cas0303 | DS40M4, ΔDSB67__07165, +pCS38 | This study |
| cas0304 | DS40M4, ΔDSB67__07165, Δ*luxO,* +pCS38 | This study |
| JAF78 | BB120, Δ*luxO*::CM | (Freeman *et al.*, 2000) |
| KT282 | BB120, *Δqrr1-5* | (Rutherford and Bassler, 2012) |
| cas0356 | BB120, Δ*luxO*::CM, +pBB147 | This study |
| cas0357 | BB120, Δ*luxO*::CM, +pLAFR2 | This study |
| KM669::pKM699 | BB120, Δ*luxR,* +pKM699 | This study |
| KM669::pLAFR2 | BB120, Δ*luxR,* +pLAFR2 | This study |
| KM83 | BB120, *luxO* D47E::CM | (Tu and Bassler, 2007) |
| cas0353 | DS40M4, *ΔluxO*:: SpecR, *ΔluxB*::TmR + *luxO,* +pCS38 | This study |
| cas0352 | DS40M4, *ΔluxR*:: SpecR, *ΔluxB*::TmR + *luxR,* +pCS38 | This study |
| cas0322 | DS40M4, Δ*cqsS,* Δ*luxPQ,* Δ*luxB*::*tm*$^{R}$ +pCS38 | This study |
| TL25 | BB120, Δ*cqsS,* Δ*luxPQ,* Δ*luxM* | (van Kessel *et al.*, 2013) |
| KT234 | BB120, *qrr1*+ | (Tu and Bassler, 2007) |
| KT212 | BB120, *qrr2*+ | (Tu and Bassler, 2007) |

| | | |
|---|---|---|
| KT225 | BB120, *qrr3*+ | (Tu and Bassler, 2007) |
| KT215 | BB120, *qrr4*+ | (Tu and Bassler, 2007) |
| KT133 | BB120, *qrr5*+ | (Tu and Bassler, 2007) |
| KT220 | BB120, *Δqrr1-5* | (Tu and Bassler, 2007) |
| KYC55 | *Agrobacterium tumefaciens* KYC55 +pJZ372 +pJZ384 +pJZ410 | (Zhu *et al.*, 2003) |
| MJ1 | | (Ruby and Nealson, 1976) |
| cas399 | DS40M4, Δ*cqsS,* Δ*luxPQ,* Δ*luxN,* Δ*CqsR*, Δ*HqsK*, Δ*luxB*::*tm*$^{R}$ , +pCS38 | This study |
| cas0408 | DS40M4, ΔCqsA, ΔluxB::SpecR, + pCS47 | This study |
| cas0409 | DS40M4, ΔluxS, ΔluxB::TmR + pCS48 | This study |
| BDP158 | DS40M4 ΔluxS, ΔcqsA, ΔluxN, ΔluxB::-SpecR + pBP15 | This study |
| BDP159 | DS40M4 ΔluxN, ΔluxB::-SpecR + BP15 | This study |
| CAS166 | DS40M4 + pCS27 | This study |

**Table S3. Plasmids used in this study.**

| Name | Description | Reference |
|---|---|---|
| pMMB67EH-tfoX-kanR | *kan*$^{R}$ , P*tac-tfoX*, *lacI* | (Simpson *et al.*, 2019) |
| pCS38 | pMMB67EH, P*luxC-luxCDABE* | This study |
| pBB147 | *qrr1*, *luxO* fragment in pLAFR2 | This study |
| pKM699 | *luxR* 2.3 kbp fragment in pLAFR2 | (Hustmyer *et al.*, 2018) |
| pLAFR2 | *V. campbellii* compatible cosmid; *tetR* | (Hustmyer *et al.*, 2018) |
| pCS47 | pMMB67EH, P*tac-cqsA* | This study |
| pCS48 | pMMB67EH, P*tac*-l*uxS* | This study |
| pBP15 | pMMB67EH, P*tac-luxN* | This study |
| pMMB67EH-kanR | pMMB67EH, empty vector | (Simpson *et al.*, 2019) |

**Table S4. Oligonucleotides used in this study.**

| Name | Sequence | Notes |
|---|---|---|
| **Primers for MuGENT or SOE products** | | |
| CAS0148 | CGTGCTCAAGTCTTCACTGATGATG | DS40M4 Δ*luxB* F1 |
| CAS0149 | GTCGACGGATCCCCGGAATGATGACTTGAT CAGAAGAACGCTTTGA | DS40M4 Δ*luxB* R1 |
| CAS0150 | GAAGCAGCTCCAGCCTACACACTCGTAACG TTTAAACGATGCTGAG | DS40M4 Δ*luxB* F2 |
| CAS0151 | GGTGAATGGCCACAAGGTACCT | DS40M4 Δ*luxB* R2 |
| CAS0303 | CTAAGCTAGAGCTGAGCGATCTCTC | F1 to make Δ*aphA* DS40M4 |
| CAS0304 | GCTAATTCAGTTTAAGCGGCCATTGGTAATG ACATGTCTTCAATCC | R1 to make Δ*aphA* DS40M4 |
| CAS0305 | ATGGCCGCTTAAACTGAATTAGCCTTGAAGT GATCGGCTAATTCGTCAC | F2 to make Δ*aphA* DS40M4 |
| CAS0306 | GTTCAGACTACTCAGCAACCGTG | R2 to make Δ*aphA* DS40M4 |
| CAS0308 | CAATGCAACTGGCATTGTTGCGAC | F1 to make Δ*qrr1* DS40M4 |
| CAS0309 | GCTAATTCAGTTTAAGCGGCCATACAGCATG GTTTGTGCCATAATTTAACC | R1 to make Δ*qrr1* DS40M4 |
| CAS0310 | ATGGCCGCTTAAACTGAATTAGCTTTGTCTG CAGATTTGGGCGCG | F2 to make Δ*qrr1* DS40M4 |
| CAS0311 | GTTGCCGCAGTTCTAATGGATAC | R2 to make Δ*qrr1* DS40M4 |
| CAS0313 | CGCATCTTCATCATATCGCCTTC | F1 to make Δ*qrr2* DS40M4 |
| CAS0314 | GCTAATTCAGTTTAAGCGGCCATAGAATTAT GCATTAATCATGCCAGTTTG | R1 to make Δ*qrr2* DS40M4 |
| CAS0315 | ATGGCCGCTTAAACTGAATTAGCTGTAGTAT ATGTACAACGTTTCCTACG | F2 to make Δ*qrr2* DS40M4 |
| CAS0316 | CAACTCTCGAGAATTAGGAAGTGG | R2 to make Δ*qrr2* DS40M4 |
| CAS0318 | CGCATCTCAAGATTCAGCAAGTTC | F1 to make Δ*qrr3* DS40M4 |
| CAS0319 | GCTAATTCAGTTTAAGCGGCCATTCACACTA AATAATGCAGTTAGTGTG | R1 to make Δ*qrr3* DS40M4 |
| CAS0320 | ATGGCCGCTTAAACTGAATTAGCTTTTATCT GGCCCCTTCAAATTCC | F2 to make Δ*qrr3* DS40M4 |
| CAS0321 | CACTTGCTCTTGTTCACCAAGGTG | R2 to make Δ*qrr3* DS40M4 |
| CAS0323 | CAGTAGACTTGGTTCGCCAATTG | F1 to make Δ*qrr4* DS40M4 |
| CAS0324 | GCTAATTCAGTTTAAGCGGCCATGTCAATCT GTATAAAGCACGATG | R1 to make Δ*qrr4* DS40M4 |
| CAS0325 | ATGGCCGCTTAAACTGAATTAGCTCTAGAG GTCAAGAATTTGCACCTC | F2 to make Δ*qrr4* DS40M4 |
| CAS0326 | GAAGAAGTGATGAATGTGGTTCGT | R2 to make Δ*qrr4* DS40M4 |
| CAS0328 | CTTGCTGGTTGGTTAGGTGTTCC | F1 to make Δ*qrr5* DS40M4 |
| CAS0329 | GCTAATTCAGTTTAAGCGGCCATGTCGCTAT AACTAAAGCATTAGGC | R1 to make Δ*qrr5* DS40M4 |
| CAS0330 | ATGGCCGCTTAAACTGAATTAGCTTTCACAA TCAATTCGCGTTTTAATTG | F2 to make Δ*qrr5* DS40M4 |

| | | |
|---|---|---|
| CAS0331 | GTAATGCTAAGCTGCGAACTTC | R2 to make Δ*qrr5* DS40M4 |
| CAS0295 | GCTAATTCAGTTTAAGCGGCCATAGGTCTCT TTGCAATTGAGTCCAT | R1 to make Δ*luxR* DS40M4 |
| CAS0296 | ATGGCCGCTTAAACTGAATTAGCATCTACAA CCGTGAACATCACTAA | F2 to make Δ*luxR* DS40M4 |
| CAS0297 | GCTAATTCAGTTTAAGCGGCCATTACCATTA GTAGATAACGAGAC | R1 to make Δ*luxO* DS40M4 |
| CAS0298 | ATGGCCGCTTAAACTGAATTAGCGTATGAAT ACGGACGTATTAAATCAGC | F2 to make Δ*luxO* DS40M4 |
| BP0338 | GTAGTCCTTGCCAGTAAAGGGCTAG | F1 to make *luxO* D47E |
| BP0339 | TCAGGTAGACGAAGCTCGAGCAGAATAAGA TCAGGAATGCGATGGTTCAGGC | R1 to make *luxO* D47E |
| BP0340 | TCTTATTCTGCTCGAGCTTCGTCTACCTGAT ATGACGGGGATGGACGTATTGC | F2 to make *luxO* D47E |
| BP0341 | TTACGACACGTTTGATCTCAGGGAG | R2 to make *luxO* D47E |
| BP0269 | ACCTCCCAAGAGACTTGGTCTACATC | F1 to make Δ*cqsA* |
| BP0270 | GCTAATTCAGTTTAAGCGGCCATCATAATAA GTTCTCTTCGTAACACAGGAAAACGC | R1 to make Δ*cqsA* |
| BP0271 | ATGGCCGCTTAAACTGAATTAGCGCCTAAC GAATCGAAAGCGGCAC | F2 to make Δ*cqsA* |
| BP0272 | CTACCTTGCGTGCGCCTTCAAC | R2 to make Δ*cqsA* |
| BP0289 | GTATGCCTTACGATGCTGCGGG | F1 to make Δ*luxS* |
| BP0290 | GCTAATTCAGTTTAAGCGGCCATCATTACAT CTCTCCTGATTGTAATCACCAACAC | R1 to make Δ*luxS* |
| BP0291 | ATGGCCGCTTAAACTGAATTAGCATCGACTA ACTTAGTTTTTAGTTTCGACAGCTG | F2 to make Δ*luxS* |
| BP0292 | GTCGACACCGATGTCTGATGCTAGG | R2 to make Δ*luxS* |
| CAS0075 | GTCGACGGATCCCCGGAATTACCATTAGTA GATAACGAGAC | R1 DS40M4 Δ*luxO* marked |
| CAS0076 | GAAGCAGCTCCAGCCTACAGTATGAATACG GACGTATTAAATCAGC | F2 DS40M4 Δ*luxO* marked |
| CAS0074 | GTCCGTCGCTCTACCAACTGAGC | F1 DS40M4 Δ*luxO* |
| CAS0079 | GCTTGTTCAACTAGGTAGCCACCAGA | R2 DS40M4 Δ*luxO* |
| CAS0084 | CCACGCAGCTCCTTGTCCGAATCG | F1 DS40M4 Δ*luxR* |
| CAS0070 | GTCGACGGATCCCCGGAATAGGTCTCTTTG CAATTGAGTCCAT | R1 DS40M4 Δ*luxR* marked |
| CAS0071 | GAAGCAGCTCCAGCCTACAATCTACAACCG TGAACATCACTAA | F2 DS40M4 Δ*luxR* marked |
| CAS0072 | CGACTACGCACACGCATCTCCAG | R2 DS40M4 Δ*luxR* |
| BP274 | CGGCTTACTCTCAAGTTTCTGTTTCGC | F1 DS40M4 Δ*CqsS* |
| BP275 | gctaattcagtttaagcggccatCGCGTCCATATTTACT TTATCTACTGAGCC | R1 DS40M4 Δ*CqsS* |
| BP276 | atggccgcttaaactgaattagcTAGTTCTTGCTCAAT AGGGTGACGAAAAC | F2 DS40M4 Δ*CqsS* |

| | | |
|---|---|---|
| BP277 | CCCAGCTCCTAACAACTGTTGTATCAC | R2 DS40M4 Δ*CqsS* |
| BP284 | ACCACATTCAAATCGTGCCATTCAAAG | F1 DS40M4 Δ*luxP/Q* |
| BP285 | gctaattcagtttaagcggccatAGAAATCAGGGAAAATAGTAACGCTTTCTTCATC | R1 DS40M4 Δ*luxP/Q* |
| BP286 | atggccgcttaaactgaattagcACCTAACGGTTAAATGGCAGTGATAAGTTAG | F2 DS40M4 Δ*luxP/Q* |
| BP287 | ATAACTCCGCTCACTCAACAGCTTAC | R2 DS40M4 Δ*luxP/Q* |
| CAS0431 | cactcccttctggctccaacattc | F1 DS40M4 ΔDSB67_07165 |
| CAS0432 | GCTAATTCAGTTTAAGCGGCCATccttggtctcataatcctttcccaac | R1 DS40M4 ΔDSB67_07165 |
| CAS0433 | ATGGCCGCTTAAACTGAATTAGCgcttgggaagcggatgaagct | F2 DS40M4 ΔDSB67_07165 |
| CAS0434 | gtagcgctctacggcaggtctc | R2 DS40M4 ΔDSB67_07165 |
| BP279 | CACTGCGTATGGCTCGTGAGAATC | F1 DS40M4 Δ*luxN* |
| BP280 | gctaattcagtttaagcggccatTGTCCATATTAGGACCAGAAAAGACACTAG | R1 DS40M4 Δ*luxN* |
| BP281 | atggccgcttaaactgaattagcTAGTTGGCACCTGATATGCATTGTTG | F2 DS40M4 Δ*luxN* |
| BP282 | GTCAGACCTACAGGATGCAGGTATCAATATC | R2 DS40M4 Δ*luxN* |
| CAS0255 | GAAGCAGCTCCAGCCTACAcaacagttggagaaggagatcagtc | F amplify DS40M4 luxO complement |
| CAS0256 | tcgtttaaacgttacgagtgcatacgttttgtttttcgtccttgc | R amplify DS40M4 luxO complement |
| CAS0252 | GAAGCAGCTCCAGCCTACAataaagtcgacttggtgagtcagtc | F amplify DS40M4 luxR complement |
| CAS0253 | tcgtttaaacgttacgagtgttagtgatgttcacggttgtagatgc | R amplify DS40M4 luxR complement |
| CAS0254 | cactcgtaacgtttaaacgatgctg | F2 DS40M4 for *ΔluxB*::complements |
| **Primers for plasmid construction** | | |
| CAS494 | GAATTCTGTTTCCTGTGTGAAATTG | R to amplify pMMB backbone |
| CAS495 | GGATCCTCTAGAGTCGACCTGC | F to amplify pMMB backbone |
| CAS496 | CAATTTCACACAGGAAACAGAATTCATGAGTG[illegible]AACC | F to amplify CqsA insert |
| CAS497 | GCAGGTCGACTCTAGAGGATCCTTAGGCGAATTCTAGTTCTGGGT | R to amplify CqsA insert |

| | | |
|---|---|---|
| CAS498 | CAATTTCACACAGGAAACAGAATTCATGCCT TTATTAGACAGCTTTACC | F to amplify LuxS insert |
| CAS499 | GCAGGTCGACTCTAGAGGATCCTTAGTCGATACGTAGCTCTTTCAG | R to amplify LuxS insert |
| BP656 | CAATTTCACACAGGAAACAGAATTCatgactga aaactttattacacacatgc | F to amplify LuxN insert |
| BP657 | AGGTCGACTCTAGAGGATCCctattctctctcagctt cacaagcg | R to amplify LuxN insert |
| **Detection Primers** | | |
| CAS0152 | ATGGCCGCTTAAACTGAATTAGC | Universal F for all unselected mutations |
| CAS0083 | GAAGCAGCTCCAGCCTACA | Universal F for all selected mutations |
| CAS0220 | GCTGAGCATCAATCGCTCTTGAC | R Δ*luxB* detect (~1kb product) |
| CAS0312 | GTACCAACACGCTCAAGATCTTG | R detect for Δ*qrr1* unmarked DS4 (~216bp) |
| CAS0317 | GAATGTACCGGAATTGGTTCGTAC | R detect for Δ*qrr2* unmarked DS4 (~317bp) |
| CAS0322 | CGCGTATTATGGTGATGTATATCTC | R detect for Δ*qrr3* unmarked DS4 (~508bp) |
| CAS0327 | CGAACTCCGAGCTTAACTCACAAG | R detect for Δ*qrr4* unmarked DS4 (~785bp) |
| CAS0332 | CCTTGGTGTTGACTATGCGATGC | R detect for Δ*qrr5* unmarked DS4 (~918bp) |
| CAS0307 | GCAAGCCAAGCGATTGATATGG | R detect *ΔaphA* unmarked DS40M4 |
| BP0273 | CGGATAAGAACAGGTTGTTCGACAAGATTG | R detect Δ*cqsA* unmarked (~150bp product) |
| BP0342 | CGAGTGATTGTGTCTACGACCATAGC | F detect *luxO* D47E (485bp upstream) |
| BP0349 | GCCAACATTCGTGATATCAACAAAGGC | R detect Δ*luxS* (~350bp product) |
| CAS0080 | CGTAGTCACGGAGTCATTGGCTTC | R Detection oligo for DS4M04 Δ*luxO* |
| CAS0085 | GAAGGCTCAATCACTGACCTTCC | R Detection oligo for DS4M04 Δ*luxR* |
| CAS0435 | ctaggttggtgctcggctatcg | R Detection oligo for DS40M4 ΔDSB67_07165 |
| BP278 | GCTGTATTCTGTCGTCCTGCAACG | R detect DS40M4 Δ*CqsS* (~250bp) |
| BP288 | GCTACGAAGCCAGTAAGAGGGC | R detect DS40M4 Δ*luxP/Q* (~450bp) |
| BP283 | GAGAGGGTAGAGGTTACGCAACGC | R detect DS40M4 Δ*luxN* (~350bp) |
| **qRT-PCR Primers** | | |
| BP345 | ATGGCTAAGGGGCAATCTCTAC | qRT-PCR DS40M4 Hfq F |
| BP346 | CTTGCAGTTTGATACCGTTCAC | qRT-PCR DS40M4 Hfq R |
| BP362 | TGGAAATCGCTCTTGAAGTGT | qRT-PCR DS40M4 luxR F |

| | | |
|---|---|---|
| BP363 | TTAAATACCGTCGCAACAGAAAC | qRT-PCR DS40M4 luxR R |
| BP364 | GTGAAATCTAGAACGCTTATTGGG | qRT-PCR DS40M4 exsB F |
| BP365 | CTTGGGAAGAGCTCGCAAAG | qRT-PCR DS40M4 exsB R |
| BP366 | ATGAAAAAGCTGCATTGGCG | qRT-PCR DS40M4 exsD F |
| BP367 | AACCTTCGTATCTTGCGCCT | qRT-PCR DS40M4 exsD R |
| BP368 | ATGTCATTACCACACGTAATTCTAACTG | qRT-PCR DS40M4 aphA F |
| BP369 | TCCAGAAGTAACCGATGCTAGC | qRT-PCR DS40M4aphA R |
| BP370 | ATGTCTGCAGAAGCACAAGC | qRT-PCR DS40M4 tssC F |
| BP371 | CACTTGGCTTTAAACGAGTTTCG | qRT-PCR DS40M4 tssC R |

### References

Bassler, B.L., Greenberg, E.P., and Stevens, A.M. (1997) Cross-species induction of luminescence in the quorum-sensing bacterium Vibrio harveyi. *J Bacteriol* **179**: 4043 LP – 4045.

Bassler, B.L., Wright, M., Showalter, R.E., and Silverman, M.R. (1993) Intercellular signalling in Vibrio harveyi: sequence and function of genes regulating expression of luminescence. *Mol Microbiol* **9**: 773–786.

Dias, G.M., Thompson, C.C., Fishman, B., Naka, H., Haygood, M.G., Crosa, J.H., and Thompson, F.L. (2012) Genome sequence of the marine bacterium Vibrio campbellii DS40M4, isolated from open ocean water. *J Bacteriol* **194**: 904.

Freeman, J.A., Lilley, B.N., and Bassler, B.L. (2000) A genetic analysis of the functions of LuxN: A two-component hybrid sensor kinase that regulates quorum sensing in Vibrio harveyi. *Mol Microbiol* **35**: 139–149.

Henke, J. and Bassler, B. (2004) Quorum sensing regulates type III secretion in Vibrio harveyi and Vibrio parahaemolyticus. *J Bacteriol* **186**: 3794–3805.

Hustmyer, C.M., Simpson, C.A., Olney, S.G., Rusch, D.B., Bochman, M.L., and van Kessel, J.C. (2018) Promoter Boundaries for the luxCDABE and betIBA-proXWV Operons in Vibrio harveyi Defined by the Method Rapid Arbitrary PCR Insertion Libraries (RAIL). *J Bacteriol* **200**: e00724-17.

van Kessel, J.C., Rutherford, S.T., Shao, Y., Utria, A.F., and Bassler, B.L. (2013) Individual and combined roles of the master regulators apha and luxr in control of the Vibrio harveyi quorum-sensing regulon. *J Bacteriol* **195**: 436–443.

de Lorenzo, V. and Timmis, K.N. (1994) Analysis and construction of stable phenotypes in gram-negative bacteria with Tn5- and Tn10-derived minitransposons. *Methods Enzymol* **235**: 386–405.

Pompeani, A.J., Irgon, J.J., Berger, M.F., Bulyk, M.L., Wingreen, N.S., and Bassler, B.L. (2008) The Vibrio harveyi master quorum-sensing regulator, LuxR, a TetR-type protein is both an activator and a repressor: DNA recognition and binding specificity at target promoters. *Mol Microbiol* **70**: 76–88.

Ruby, E.G. and Nealson, K.H. (1976) Symbiotic association of Photobacterium fischeri with the marine luminous fish  Monocentris japonica; a model of symbiosis based on bacterial studies. *Biol Bull* **151**: 574–586.

Rutherford, S.T. and Bassler, B.L. (2012) Bacterial quorum sensing: Its role in virulence and possibilities for its control. *Cold Spring Harb Perspect Med* **2**: 1–25.

Simpson, C.A., Podicheti, R., Rusch, D.B., Dalia, A.B., and van Kessel, J.C. (2019) Diversity in Natural Transformation Frequencies and Regulation across Vibrio Species. *MBio* **10**: 1–16.

Tu, K.C. and Bassler, B.L. (2007) Multiple small RNAs act additively to integrate sensory information and control quorum sensing in Vibrio harveyi. *Genes Dev* **21**: 221–233.

Waters, C.M. and Bassler, B.L. (2006) The Vibrio harveyi quorum-sensing system uses shared regulatory components to discriminate between multiple autoinducers. *Genes Dev* **20**: 2754–2767.

Zhu, J., Chai, Y., Zhong, Z., Li, S., and Winans, S.C. (2003) Agrobacterium Bioassay Strain for Ultrasensitive Detection of N-Acylhomoserine Lactone-Type Quorum-Sensing Molecules: Detection of Autoinducers in Mesorhizobium huakuii. *Appl Environ Microbiol* **69**: 6949–6953.
